# Phenotypic Heterogeneity Facilitates Survival While Hindering the Evolution of Drug Resistance Due to Intraspecific Competition

**DOI:** 10.1101/2021.09.20.460867

**Authors:** Joshua Guthrie, Daniel A. Charlebois

## Abstract

Rising rates of resistance to antimicrobial drugs threatens the effective treatment of infections across the globe. Drug resistance has been established to emerge from non-genetic mechanisms, such as “persistence” in quiescent microbes and fluctuations in gene expression in actively replicating cells, as well as from genetic mutations. However, it is still unclear how non-genetic drug resistance affects the evolution of genetic drug resistance. We develop deterministic and stochastic population models that incorporate resource competition to quantitatively investigate the transition from non-genetic to genetic resistance during the exposure to static and cidal drugs. We find that non-genetic resistance facilitates the survival of cell populations during drug treatment, but that it hinders the development of genetic resistance due to the competition between the non-genetically and genetically resistant subpopulations. Non-genetic drug resistance in the presence of subpopulation competition is found to increase the first-appearance and fixation times of drug resistance mutations, while increasing the probability of mutation before population extinction during cidal drug treatment. Intense intraspecific competition during drug treatment leads to extinction of the susceptible and non-genetically resistant subpopulations. These findings advance our fundamental understanding of the evolution of drug resistance and may guide novel treatment strategies for patients with drug-resistant infections.

**SIGNIFICANCE:** Drug resistance is predicted to kill as many as 10 million people per year and cost over 100 trillion USD in cumulative lost production globally by 2050. To mitigate these socio-economic costs, we need to fundamentally understand the drug resistance process. We investigate the effect that different forms of resistance have on the evolution of drug resistance using mathematical modeling and computer simulations. We find that the presence of non-genetically drug-resistant cells (whose resistance is temporary and not encoded in a genetic mutation) allows the population to survive drug treatment, while competition between these subpopoulations simultaneously slows down the evolution of permanent genetic drug resistance and in some cases drives them extinct. These findings have important implications for advancing evolutionary theory and for developing effective “resistance-proof” treatments.

## INTRODUCTION

Antimicrobial (drug) resistance occurs when bacteria, viruses, fungi, and parasites no longer respond to drug therapy, making infections difficult or impossible to treat, which increases the risk of disease transmission, severe illness, and death (1). The evolution of drug resistance is well known to arise from the natural selection of genetic (DNA or RNA) mutations that provide microbes with the ability to survive and proliferate during treatment (2). More recently, it has been established that non-genetic mechanisms promote microbial phenotypic diversification and survival strategies in selective drug environments (3, 4). Phenotypic heterogeneity has important implications for drug resistance (3, 5), with heritable resistance potentially arising independently of genetic mechanisms (6). The stochastic or “noisy” expression of genes (7, 8) introduces phenotypic variation among genetically identical cells in the same drug environment, which can result in the fractional killing of clonal microbial populations (4, 9) and chemotherapy resistance in cancer (10). This stochasticity is due in part to the inherently random nature of the biochemical reactions involved in the transcription and translation of genetic material, and can lead to the emergence of phenotypically distinct subpopulations within an actively dividing clonal cell population (8, 11). Another form of non-genetic drug resistance occurs in non-growing or slow-growing bacterial “persister” cells, which generate the phenotypic variation required for clonal bacterial populations to survive antibiotics that require active growth for killing (12–14); a similar phenomenon is now thought to occur in fungi (15) and cancer (16).

Non-genetic drug resistance (temporary resistance that is not encoded in a genetic mutation) has been proposed to promote the development of genetic drug resistance (permanent resistance that is acquired through drug resistance mutations or genes) (3, 5, 10, 17). This process may be enhanced by the interaction between non-genetic and genetic mechanisms inside the cell (5). For instance, promoter mutations can alter the expression noise levels of drug resistance genes (9), genetic networks can modulate gene expression noise to enhance drug resistance (18–20), and stress response genes can evolve elevated transcriptional variability through natural selection (21, 22). Non-genetic mechanisms can facilitate the generation of genetic diversity by increasing the expression of key regulators involved in DNA replication, recombination, or repair (23, 24), as well as by enhancing the adaptive value of beneficial mutations during drug treatment (25) and promoting the fixation of favorable gene expression altering mutations (26). However, there are conflicting views on how phenotypic heterogeneity may facilitate adaptive evolution (27) and the transition from non-genetic to genetic drug resistance remains to be quantified (5). Importantly, the interactions between non-genetic and genetic resistance mechanisms have not been investigated in the context of resource competition between these subpopulations in a microbial population undergoing drug treatment.

Non-genetic phenotypic variability can impact cellular population dynamics by providing a link between micro-scale dynamics (such as stochasticity at the molecular level) and macro-scale biological phenomena (including the fate of interacting cell populations) (28). Such noise in biological systems may facilitate the adaption to environmental stress by allowing distinct, co-existing cellular states in a population to find the best adaptive solution from multiple starting points (29). Phenotypic heterogeneity can also promote interactions among subpopulations as well as the division of labor between individual cells, providing clonal microbial populations with new functionalities (30), which has not been investigated in the context of the interaction between non-genetic and genetic mechanisms. Finally, intraspecific competition leads to logistic population growth; population growth is exponential when population size and resource competition are low, followed by a progressively reduced growth rate as the population size increases towards the carrying capacity of the microenvironment (31).

In this study, we investigate the transition from non-genetic to genetic resistance during static drug (drugs that slow or stop cell growth) and cidal drug (drugs that kill cells) treatment in the presence of resource competition using deterministic and stochastic population models (31). Overall, we find that non-genetic resistance facilitates the initial survival of cell populations undergoing drug treatment, while hindering the fixation of genetic mutations due to competition effects between non-genetically and genetically resistant subpopulations. Quantitatively understanding the interaction between non-genetic and genetic mechanisms will improve our fundamental understanding of drug resistance evolution and may lead to more effective treatments for patients with drug-resistant infections.

## METHODS

### Deterministic Population Model

The deterministic population model describes changes in cellular subpopulation concentrations over time under the influence of a drug. Three different subpopulations comprising the total population *T* are described in this model: a susceptible subpopulation *S*, a non-genetically resistant subpopulation *N*, and a genetically resistant subpopulation *G*. Cells may switch between the *S* and *N* subpopulations, and cells in the *N* subpopulation can mutate into the *G* subpopulation (Fig. 1). The mathematical model is described by a set of coupled ordinary differential equations (ODEs):

**Figure 1:**
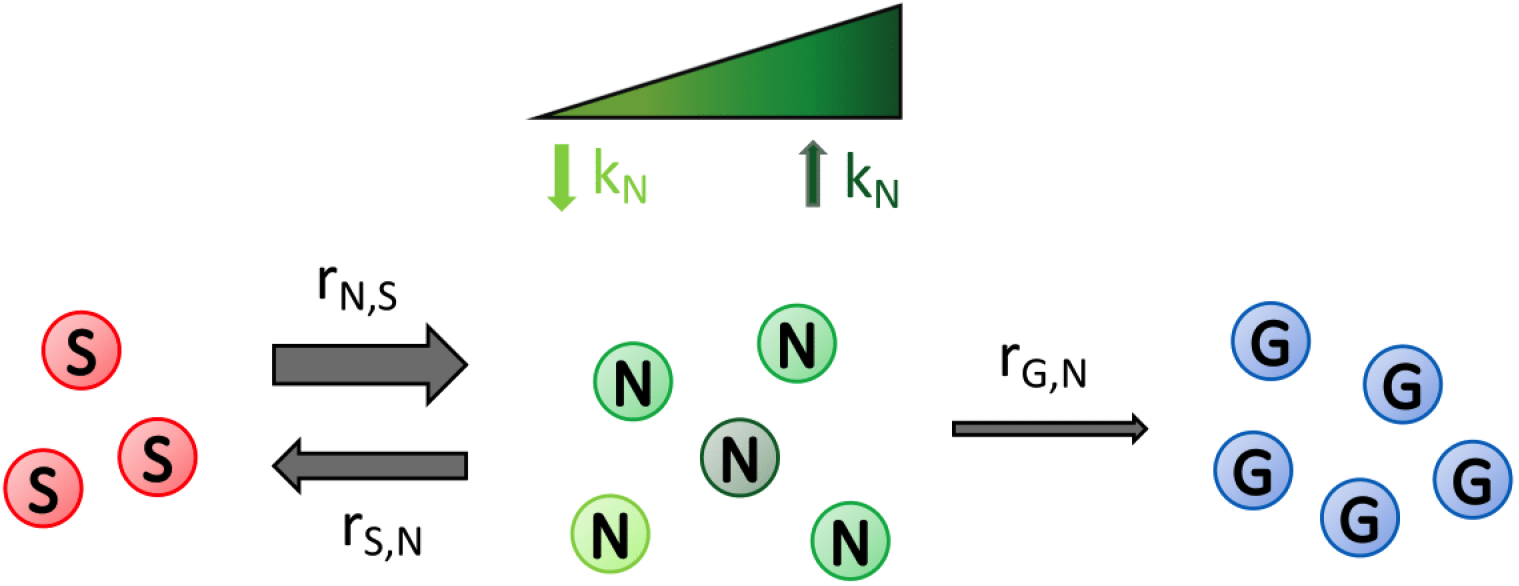
Schematic depicting the transitions between susceptible, non-genetically resistant, and genetically resistant subpopulations in a cell population undergoing drug treatment. Cells from the drug susceptible (*S*) and the non-genetically drug-resistant (*N*) subpopulations can switch between these phenotypes (at rates *r*_*N,S*_, and *r*_*S,N*_, respectively). *N* cells have varying degrees of transient drug resistance (dark green cells are more drug resistant than light green cells), as described by their fitness (growth rate, *k*) in the presence of a drug. Cells from the *N* subpopulation can mutate (at a rate *r*_*G,N*_) to become permanently genetically drug-resistant (*G*) cells.

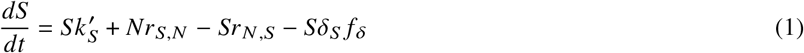

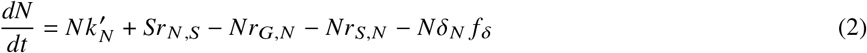

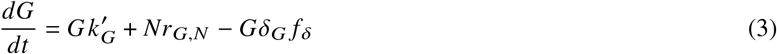

where *r*_*S,N*_, is the switching rate from *N* to *S, r*_*N,S*_, is the switching rate from *S* to *N, r*_*G,N*_ is the mutation rate from *N* to *G, δ*_*S*_, *δ*_*N*_, *δ*_*G*_ are the death rates of *S, N*, and *G*, respectively, and *f* _*δ*_ is a dimensionless death rate scaling factor. 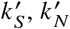 and 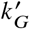 describe the growth of each subpopulation in the presence of a drug and resource competition and are given by Eq. 6. There is no mutational pathway from *S* to *G*, as we are considering drugs that completely arrest growth and division 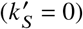 or kills (*f*_*δ*_ *>* 1) *S* cells and therefore genetic mutation due to DNA replication errors does not occur. Unless otherwise indicated, we assume that *N* has partial, temporary resistance (i.e., 0 *< δ*_*N*_ *< δ*_*S*_) and that genetic mutation provides complete, permanent resistance to the drug (*δ*_*G*_ *f* _*δ*_ = 0).

To model cidal drug treatments of varying strength we multiplied the deaths rates in Eqs. 1-3 by *f* _*8*_. Summing Eqs. 1-3 under the above assumptions yields the following equations:

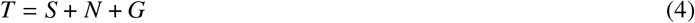

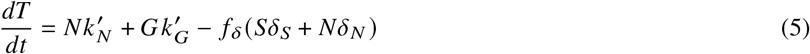

Resource competition between the subpopulations was modelled by scaling *k*_*N*_ and *k*_*G*_ by a Baranyi-Hill type function, which depends on *T* and results in logistic growth (31). For subpopulation *i* (where, *i* ∈ {*S, N, G*}), this is given by:

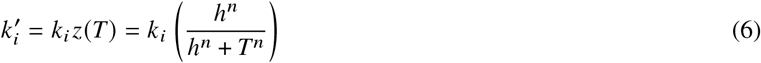

where *k*_*i*_ is the growth rate for subpopulation *i* (which leads to exponential growth in the absence of competition for limited resources), *n* is the Hill coefficient, and *h* is the point at which the competition function *z* (*T*) is half of its maximum value.

The growth dynamics of *S, N*, and *G* were obtained by solving the deterministic model, starting from initial population sizes *S*_*i*_, *N*_*i*_, and *G*_*i*_ and numerically integrating Eqs. 1-3 over a total time *t*_*tot*_ using a time step Δ_*t*_. This numerical integration was performed using the *ode45* ODE solver, which is based on an explicit Runge-Kutta method, in MATLAB (32). The steady-state solutions to Eqs. 1-3 are presented in the Supporting Material (Section 6). The fixation time *τ*_*fix*_ was used as a quantitative measure of how long it takes for *G* to become dominant in the population (33) and was defined as the time it takes for *G* to comprise 95% of the total population.

### Stochastic Population Model

Next, we developed a stochastic population model corresponding to the deterministic population model to study the effect of non-genetic resistance on the evolution of genetic resistance in low cell number regimes. Low numbers of infectious cells can occur during drug treatment and is the regime where stochastic fluctuations are expected to have a significant effect on population dynamics. Accordingly, Eqs. 1-3 were translated into the following set of reactions:

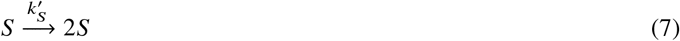

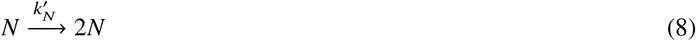

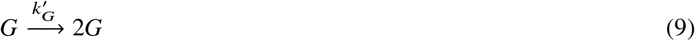

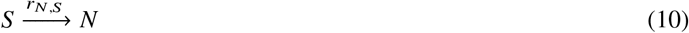

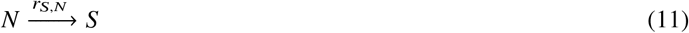

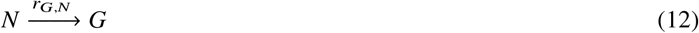

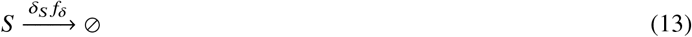

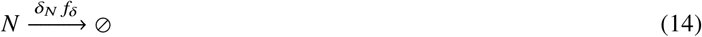

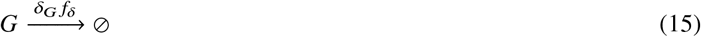

Eqs. (7)-(15) were simulated using the Gillespie stochastic simulation algorithm (34, 35).

To quantify the effect of non-genetic drug resistance on the evolution of genetic drug resistance in the stochastic population model, we obtained the first-appearance time (*P*_*τ*_) and fixation time 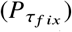 distributions of *G*. For parameter regimes where population extinction could occur during the cidal drug treatment simulations (i.e., when *S* and *N* go extinct before *G* appears), we determined the effect that the growth rate (fitness) of *N* had on the probability of *G* emerging (*P*_*G*_) before the extinction of the population. This was calculated from the number of population extinction events that occurred over a large number of simulations for different values of *k*_*N*_.

While the deterministic population model (Eqs. 1-3) was suitable for investigating large population dynamics under drug treatment, the corresponding stochastic population model (Eqs. 7-15) was necessary to accurately quantify fixation time and mutation first-appearance time distributions and extinction events for cidal drug treatment scenarios where the total population size becomes small.

## RESULTS AND DISCUSSION

The parameters for the deterministic model and stochastic population model are provided in the Supporting Material (Table S1) and the simulation codes are freely available at: https://github.com/CharleboisLab/S-N-G.

### Deterministic Population and Evolutionary Dynamics Under Static Drug Exposure

We began by numerically solving the deterministic population model to generate the time series of subpopulation concentrations to investigate the relative fitness effects of the non-genetically resistant and genetically resistant subpopulations on the evolution of drug resistance during static drug treatment.

The concentration of *S* initially decreases after the application of the static drug as a result of cells switching from *S* to *N*, and then increases logistically due to switching from *N* to *S* (Fig. 2A). The growth of *N* follows a logistic-type curve (Fig. 2B). Overall, the growth of *S* and *N* (Fig. 2A,B), along with the growth of total population (Fig. 2D), increase as the fitness of *N* increases.

**Figure 2:**
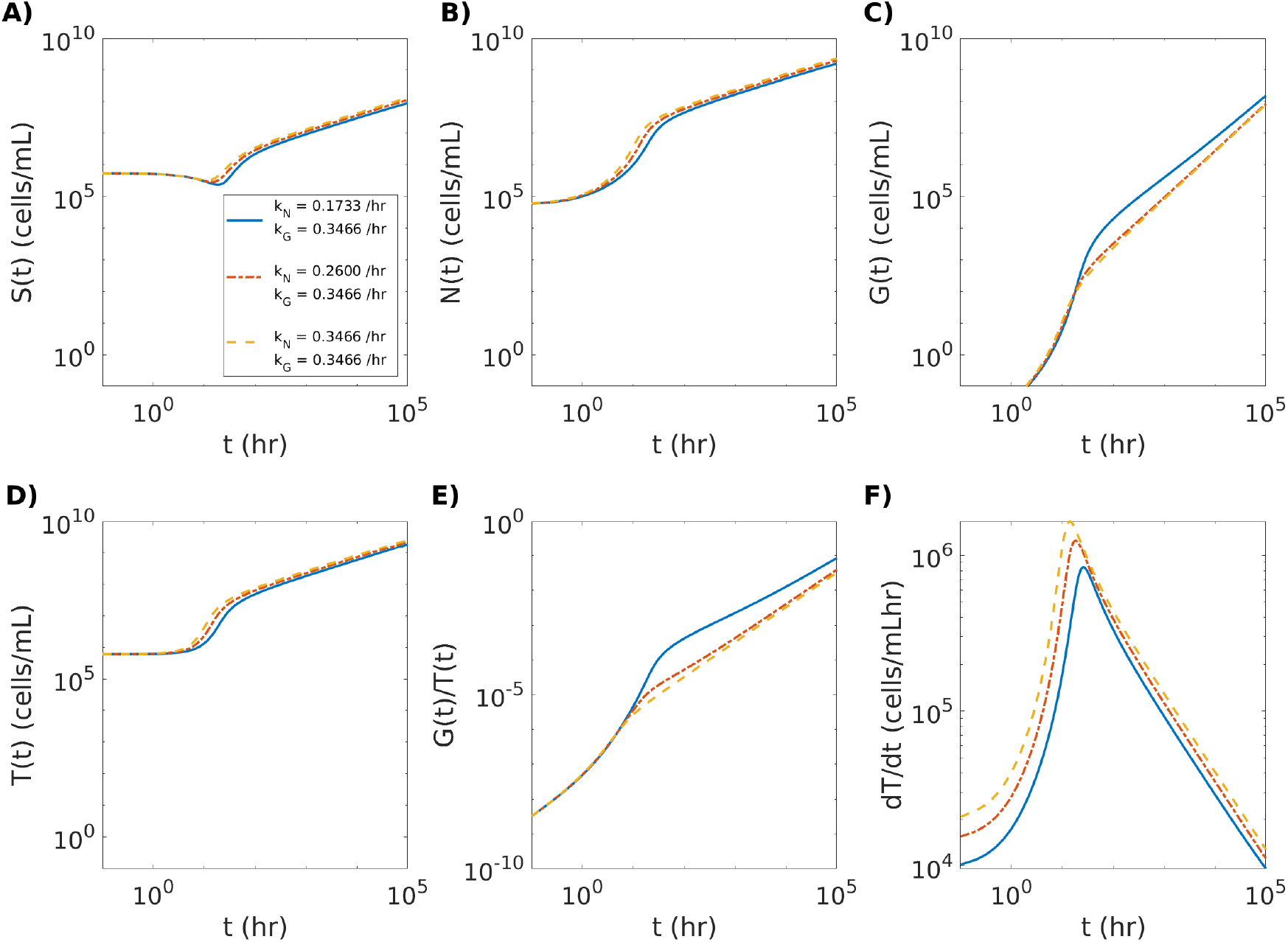
Growth of the genetically resistant subpopulation is hindered by an increase in growth rate of the non-genetically resistant subpopulation in a static drug environment. (A) The growth curve of the susceptible (*S*) subpopulation. (B) The growth curve of the non-genetically resistant (*N*) subpopulation. (C) The growth curve of the genetically resistant (*G*) subpopulation. (D) The growth curve of the total population (*T*). (E) The fraction of *G* in the total population (*T*). (F) The rate of change in the size of *T* (*dT*/ *d*) as a function of time (). Each colored line represents a different numerical simulation corresponding to the growth values shown in the legend in (A), with the solid blue line representing the lowest level of *N* fitness (growth rate of 0.1733 /hr), the red dash-dotted line an intermediate level of *N* fitness (growth rate of 0.2600 /hr), and the yellow dashed line the highest level *N* fitness (growth rate of 0.3466 /hr) relative to the fitness of *G* (growth rate of 0.3466 /hr) during static drug treatment.

The concentration of *G* (Fig. 2C) and the fraction of *G* in the total population (Fig. 2E) reveal that an increase in the fitness of *N* decreases the expansion of *G*. This can be attributed to resource competition between the subpopulations, as a higher total population concentration reduces the growth rate of *G* (Eq. (6)).

The growth rate of the total population over time increases before sharply decreasing after it reaches a maximum (Fig. 2F). This is a result of the growth of the population beginning to slow down as it increases in size (*dT* /*dt* → 0 as *T*→ ∞), which is expected for logistic-type growth (Eq. (6)) (31). Despite the decrease in the expansion of *G*, the total population growth rate increases as the fitness of *N* increases (Fig. 2F). Thus, increasing the fitness of *N* in the static drug environment enhances the growth of the population, while at the same time hindering the expansion of *G*. Additionally, intense resource competition can drive the *S* and *N* subpopulations to extinction over longer timescales (Fig. S1).

Then, we quantified how the fitness of the *N* and *G* subpopulations affects the fixation of the mutated *G* subpopulation. As expected, an increase in *k*_*G*_ relative to *k*_N_ shortens the fixation time of *G* in the population (Fig. 3A). Importantly, increasing *k*_*N*_ relative to *k*_*G*_ lengthens the fixation time of *G* (Fig. 3A), due to competition decreasing the growth of *G* (Fig. 2C,E). Overall, the trends in the static drug environment were qualitatively similar for a wide range of parameters (Figs. S2-S5).

**Figure 3:**
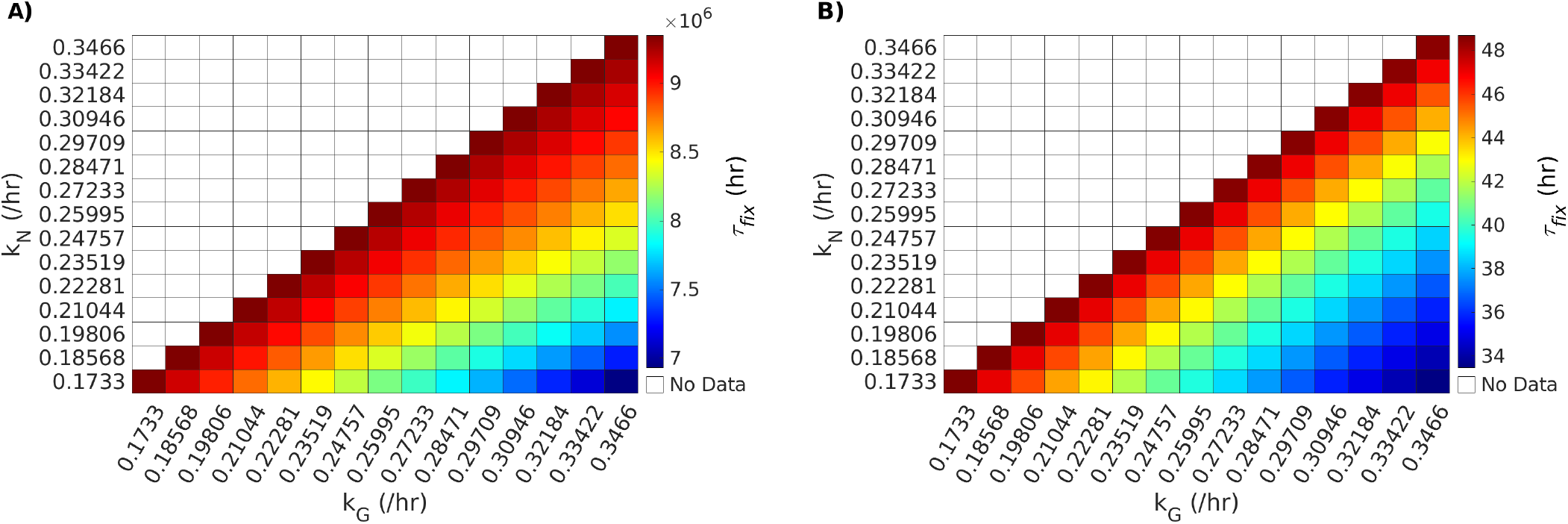
Drug resistance of the non-genetic subpopulation slows the evolution of the genetically drug resistant subpopulation during drug treatment. (A) Heat map shows the effect of the growth rates of the non-genetically resistant (*k*_*N*_) and genetically resistant (*k*_*G*_) subpopulations on the fixation time (*τ* _*fi x*_) of *G* during static drug treatment. (B) Heat map shows the effect of *k*_*N*_ and *k*_*G*_ on the *τ*_*fix*_ of *G* under cidal drug treatment. The cidal death rate scaling factor was set to *f* _*δ*_ = 2 for these simulations. Each bin in (A) and (B) corresponds to a simulation for a particular combination of *k* and *k*_*G*_ parameter values. The color map gives *τ*_*fix*_ in hours. As *k*_*N*_ is less than or equal to *k*_*G*_, numerical simulation data does not appear in the upper diagonal of the heat maps.

### Deterministic Population and Evolutionary Dynamics Under Cidal Drug Exposure

Next, we investigated how the relative fitness of the non-genetically resistant and genetically resistant subpopulations affected the evolutionary dynamics of the population under cidal drug treatment.

The concentration of *S* quickly declines after exposure to the cidal drug, with switching from *N* providing temporary survival before dying off (Fig. 4A). The *N* subpopulation shows slight temporary growth for higher values of *k*_*N*_ before dying off, while lower values of *k*_*N*_ produce flat growth curves before going extinct due to drug treatment and intraspecific competition (Fig. 4B). Higher *k*_*N*_ values prolong the temporary survival of *S* and *N* compared to lower *k* _*N*_values (Figs. 4A,B). The logistic growth of *G* was unaffected by the fitness of *N* (Fig. 4C). Increasing the fitness of *N* decreases the fraction of *G* in the total population, as higher *k*_*N*_ values result in more *N* cells and consequently lower *G*/ *T* values (Fig. 4E).

**Figure 4:**
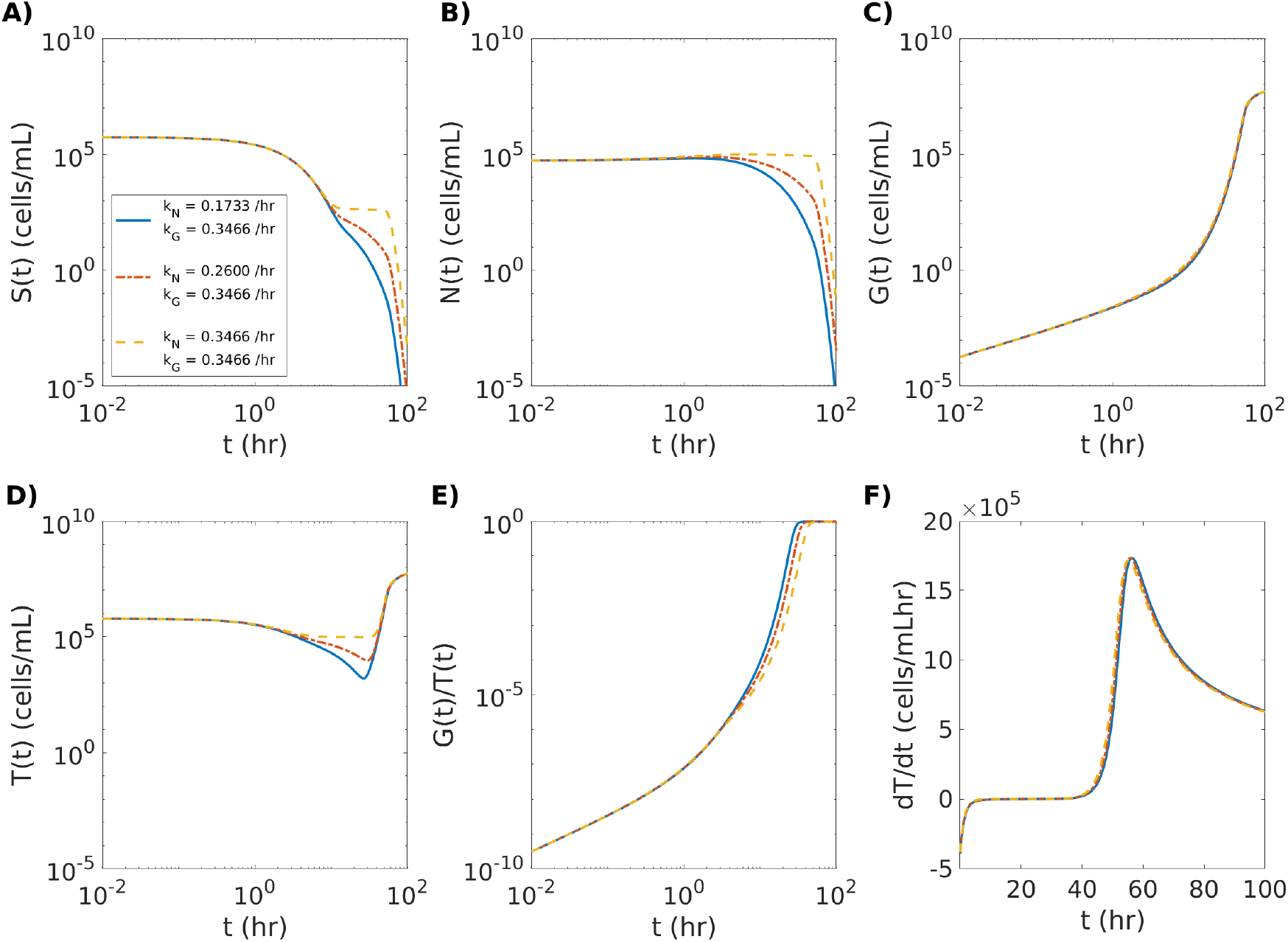
The fraction of the genetically resistant cells in the population is reduced by an increase in growth rate of the non-genetically resistant subpopulation in a cidal drug environment. (A) The growth curve of the drug susceptible (*S*) subpopulation. (B) The growth curve of the non-genetically drug resistant (*N*) subpopulation. (C) The growth curve of the genetically drug resistant (*G*) subpopulation. (D) The growth curve of the total population (*T*). (E) The fraction of *G* in the total population (*T*). (F) The rate of change in the size of *T* (*dT*/ *d*) as a function of time (*t*). Each colored line represents a different numerical simulation corresponding to the growth values shown in the legend in (A), with the solid blue line representing the lowest level of *N* fitness (growth rate of 0.1733 /hr), the red dash-dotted line an intermediate level of *N* fitness (growth rate of 0.2600 /hr), and the yellow dashed line the highest level *N* fitness (growth rate of 0.3466 /hr) relative to the fitness of *G* (growth rate of 0.3466/hr) during static drug treatment. The cidal death rate scaling factor was set to *f* _*δ*_ = 2 for these simulations.

The number of cells in the total population initially declines (negative growth rate) before stabilizing at zero population growth (Fig. 2D,F). Then the population growth rate increases as the genetically drug resistant *G* subpopulation expands to take over the population. This is followed by a decrease in the population growth rate, as the total population size moves towards saturation after the maximum population growth rate is reached. When *G* is considered to be partially resistant (i.e. *δ*_*G*_ *f* _*δ*_ *>* 0), cidal drug treatment can either drive the total population to extinction or result in non-zero steady-state *S, N*, and *G* subpopulation concentrations (Fig. S6).

As for the static drug treatment, increasing the fitness of *N* hinders the fixation time of the genetically resistant subpopulation during cidal drug treatment (Fig. 3B). Similar population dynamics were observed for weaker cidal drug (*f* _*δ*_ = 1; Fig. S7) and stronger cidal drug (*f* _*δ*_ = 4; Fig. S8) scenarios when compared to the *f* _*δ*_ = 2 scenario (Fig. 3B). We also found that by comparison the *f* _*δ*_ = 2 scenario that weaker cidal drug (Fig. S8A) and stronger cidal drug (*f* _*δ*_ = 4; Fig. S8B) scenarios had intermediate fixation times. Overall, the findings for the cidal drug environment were qualitatively the same for a wide range of parameters (Figs. S9-S13).

### Stochastic Evolutionary Dynamics During Cidal Drug Treatment

We simulated the stochastic population model corresponding to the deterministic population model to investigate the transition from non-genetic to genetic drug resistance in cell populations moving towards extinction during cidal drug treatment. This is important as fluctuations in small subpopulation sizes may impact the evolutionary dynamics of the population. Given that *S* and *N* are killed by differing degrees by the cidal drug, and that the evolution of *G* depends directly on *N*, we anticipated that stochastic fluctuations in the size of *N* would in some cases lead to the extinction of the total population. To quantify the population and evolutionary drug resistance dynamics in this regime, we determined how the fitness of *N* affects the probability of population extinction (which occurs when *S* and *N* go extinct before *G* emerges; Fig. S14), along with the first-appearance time distribution of a genetic mutation, and the fixation times which are bound to occur in our model once *G* is present in the population.

Increasing the fitness of *N* increased the likelihood of *G* appearing and rescuing the cell population from extinction during intermediate strength cidal drug treatment (*f* _*δ*_ = 2; Fig. 5A). This is in agreement with previous work that found that increasing the fluctuation relaxation time of a drug resistance gene increased the probability of acquiring a drug resistance mutation (6). When *N* had low fitness in the cidal drug (*k*_*N*_ = 0.1733 /hr) the population went extinct 83% of the time. The extinction probability decreased exponentially as the fitness of *N* increased, with the population going extinct 69% of the time for moderate fitness (*k*_*N*_ = 0.2600 /hr) and 0.00163% of the time for high fitness (*k*_*N*_ = 0.3466 /hr). These results show that the presence of non-genetic resistance enhances the survival of the population when there are no pre-existing drug resistance mutations prior to drug exposure, and that non-genetic resistance increases the chance that a mutation will occur by providing the population with more time before extinction due to drug treatment. As expected, *G* always appeared in the population when the strength of the cidal drug was low (*f* _*δ*_ = 1), and conversely, the population almost always went extinct before *G* could emerge when the strength of the cidal drug was high (*f* _*δ*_ = 4) (Fig. 5A). Decreasing the fitness of *N* resulted in an earlier but more gradual transition from population survival to population extinction as the strength of the cidal drug increased (Fig. 5B).

**Figure 5:**
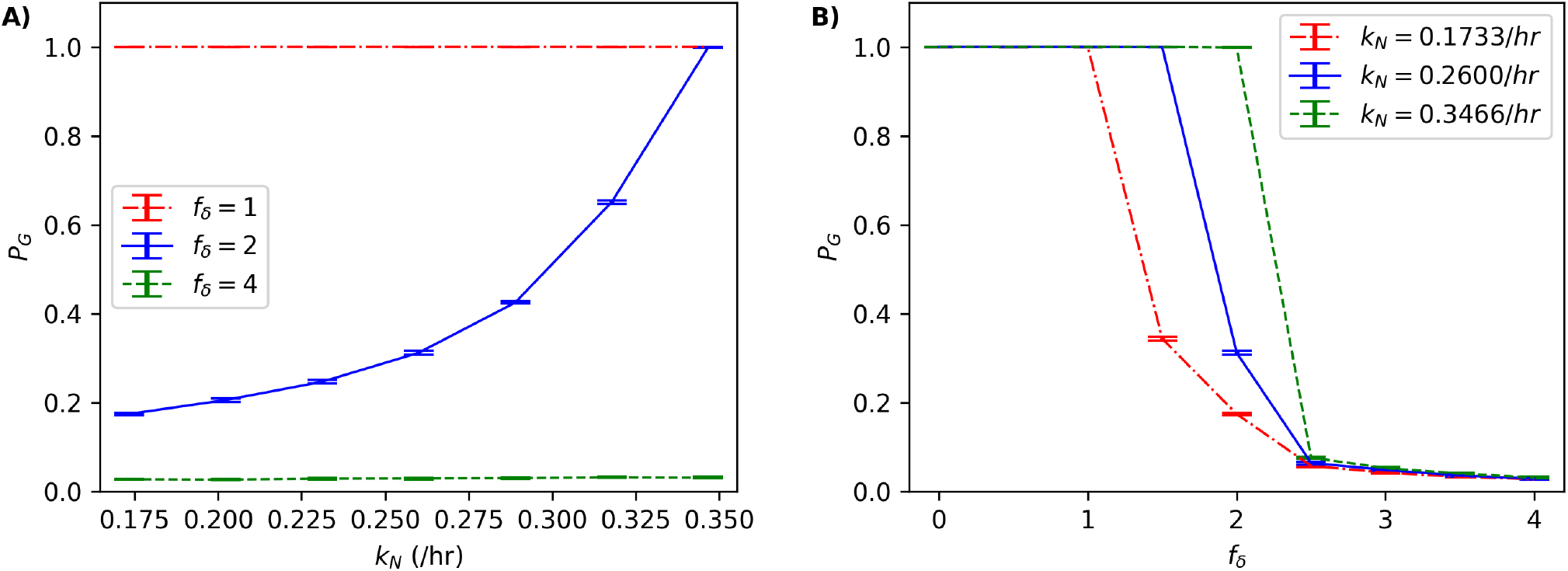
Probability of genetic drug resistance emerging before population extinction due to cidal drug treatment. (A) The appearance probability of the genetically drug resistant subpopulation (*P*_*G*_) as a function of *N* fitness (growth rate) values of non-genetic subpopulation (*k*_*N*_). Each colored line represents a different strength of the cidal drug, with the red dashed-dotted line representing the lowest drug strength (*f* _*δ*_ = 1), the blue line an intermediate drug strength (*f* _*δ*_ = 2), and the green dashed-dotted line the highest drug strength (*f* _*δ*_ = 4). (B) The appearance probability *G* as a function of cidal drug strength. Each colored line represents a different level of drug resistance for *N*, with the red dashed-dotted line representing the lowest fitness (growth rate of *k*_*N*_ = 0.1733 /hr), the blue line an intermediate fitness (growth rate of *k*_*N*_ = 0.2600 /hr), and the green dashed-dotted line the highest fitness (growth rate of *k*_*N*_ = 0.3466 /hr). Each data point in (A) and (B) is an average over ten realizations of 10, 000 simulations. Error bars show the standard deviation.

When the fitness of *N* increased in the cidal drug environment so did the means of the mutation first-appearance time and fixation time distributions (Fig. 6). This indicates that the presence of non-genetic drug resistance slows the evolution of genetic drug resistance, in agreement with the results obtained from the deterministic population model (Fig. 3B). While the coefficient of variation (CV; defined as the standard deviation divided by the mean) of the first-appearance time distributions (Fig. 6A,C,E) was only marginally dependent on the fitness of *N*, the CV of the fixation time distributions (Fig. 6B,D,F) increased approximately three fold as the fitness of *N* increased from low to high. Therefore, the presence of increased non-genetic drug resistance is predicted to not only slow down genetic drug resistance, but also to increase the uncertainty in its evolution.

**Figure 6:**
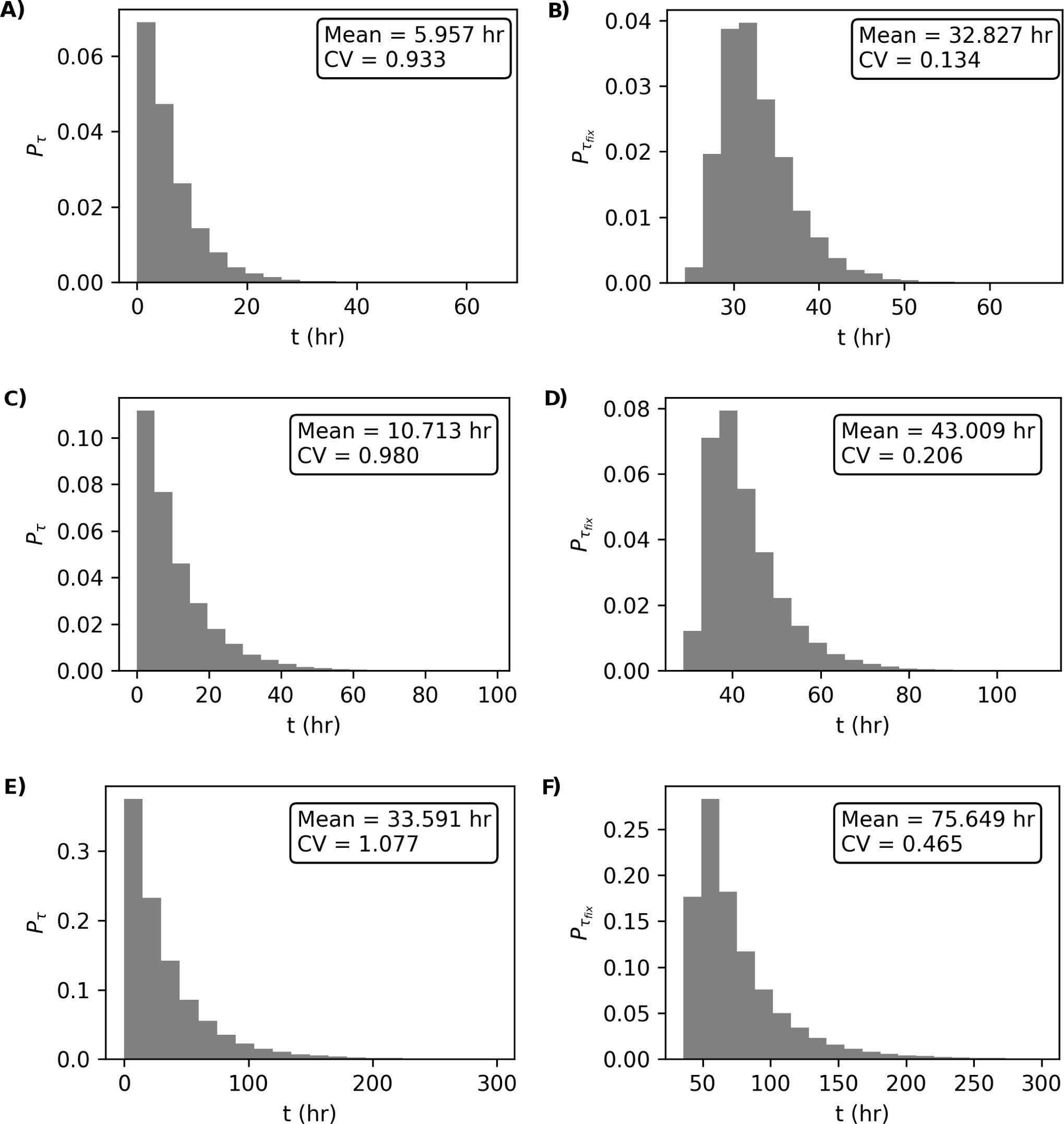
Drug resistance mutation first-appearance time and fixation time distributions during cidal drug treatment. (A) First-appearance time 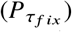 and (B) fixation time (*P*_*τfix*_) distributions for the genetically resistant subpopulation *G* for low non-genetically resistant subpopulation *N* fitness (growth rate; *k*_*N*_ = 0.1733 /hr). For (A)-(B), population extinction occurred for 82, 586 out of 100, 000 simulations. (C) *P*_*τ*_ and (D) 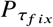 distributions for *G* for intermediate *N* fitness (*k*_*N*_ = 0.2600 /hr). For (C)-(D), population extinction occurred for 68, 812 out of 100, 000 simulations. (E) *P*_*τ*_ and (F)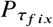 distributions for *G* for high *N* fitness (*k*_*N*_ = 0.3466 /hr). For (E)-(F), population extinction occurred for 163 out of 100, 000 simulations. The mean and the coefficient of variation (CV) for each distributions is provided in the top right-hand corner of panels (A)-(F). The cidal death rate scaling factor was set to *f* _*δ*_ = 2 for these simulations.

## CONCLUSION

We found using deterministic and stochastic population models that while non-genetic resistance enhances population survival during drug treatment, a slower rate of genetic resistance evolution emerges from resource competition between these subpopulations. More specifically, increasing the fitness of the non-genetically resistant subpopulation (which allows the population to survive initial drug exposure) exponentially increased the chance of a genetically resistant subpopulation appearing and rescuing the population from extinction during cidal drug treatment. However, increasing the fitness of the non-genetically resistant subpopulation slowed down the fixation time of genetic drug resistance mutations due to competition effects, when no pre-existing mutations were present in the population. Incorporating pre-existing mutations into the model is expected to reduce the timescale of fixation, as it would remove the growth delay resulting from the time it takes for non-genetically resistant cells to mutate into genetically resistant cells. Furthermore, high levels of intraspecific competition drove the susceptible and non-genetically resistant subpopulations extinct in static and cidal drug treatment scenarios, which opens the possibility of incorporating competition and resource limitations strategies into antimicrobial therapies. These predictions could be tested experimentally, for instance through competition assays (36) in which synthetic gene circuits tune the initial fractions of susceptible and non-genetically drug resistant cells in the population (37). Overall, our quantitative model generated robust and novel predictions on the evolution of drug resistance, and revealed that the interplay between transient non-genetic drug resistance and permanent genetic resistance may be more complex than previously thought (3, 5, 10, 17, 25). As drug exposure is more generally a form of selective pressure, the results of this study are also anticipated to be useful for advancing evolutionary theory.

It will be important to study the effects of non-genetic resistance on the development of genetic resistance in the context of fluctuating environments, which may be governed by environment-sensing genetic networks (38), along with cellular trade-offs that may occur in drug environments (39). Fluctuating environmental stressors have been shown to facilitate ‘bet-hedging’ in cell populations (40, 41), whereby some cells adopt a non-growing, stress-resistant phenotype to increase the long-term fitness of the population. This could be modeled using stochastic hybrid processes (42), for instance by using an stochastic ON-OFF switch coupled to a system of ordinary differential equations describing subpopulation dynamics in the presence of a drug. Furthermore, the first-appearance time, fixation time, and extinction events could be described analytically in future studies using a first-passage time framework (6, 43).

A complete understanding of the drug resistance process, including the interplay between non-genetic and genetic forms of drug resistance, will be important for mitigating the socio-economic costs of antimicrobial resistance (5). The drug resistance mutation appearance probabilities and first-appearance time distributions determined using stochastic population models may prove useful for guiding patient treatment. During the course of treatment, a drug could be substituted or combined with another drug (44, 45) before permanent, genetic drug resistance is predicted to emerge. Fluctuations in mutation first-appearance times are also important, as they can be the difference between the eradication or the establishment of drug-resistant infection during drug therapy. Finally, drug-resistant infections may one day be overcome by novel strategies that enhance competition between non-genetically and genetically resistant pathogens during treatment.

## AUTHOR CONTRIBUTIONS

D.C. conceived, designed, and supervised the research. D.C. developed the mathematical models. J.G. carried out the numerical and stochastic simulations. J.G. and D.C. analyzed the data. D.C. and J.G. wrote the article.

## ACKNOWLEDGMENTS

We thank Dr. Kevin S. Farquhar for helpful discussions. D.C. was financially supported by the Government of Canada’s New Frontiers in Research Fund—Exploration program (2019-01208) and the University of Alberta. J.G. was supported by a 2021 NSERC USRA.

## SUPPORTING CITATIONS

References (46–48) appear in the Supporting Material.

## Supporting Material

An online supplement to this article can be found by visiting BJ Online at http://www.biophysj.org.

### 1 Parameters

The parameter values used in our study (Table 1) were based on the literature or were scanned over a range of values.

Drug susceptible cells *S* cells formed the majority of the total initial population (*T*_*i*_) and non-genetically drug resistant cells *N* formed 1-10% of *T*_*i*_ [2]. We assumed there were no pre-existing genetic drug resistance mutations (*G*_*i*_ = 0) for all numerical and stochastic simulations. Numerical simulation of the deterministic population started with *S*_*i*_ = 5.5×10^5^ cells/mL and *N*_*i*_ = 5.5×10^4^ cells/mL, which are common initial concentrations for ‘log-phase’ laboratory experiments [1].

Depending on the concentration of the drug being considered, constant growth rates *k*_*N*_ and *k*_*G*_ (which determine the fitness of the given subpopulation in the presence of a drug) were assigned values between 0.1733 /hr and 0.3466 /hr, based on growth rates measured in standard yeast cell culture experiments [4]. These parameter ranges were the basis of parameter scans that were used to predict how the relative fitness between *N* and *G* will affect population dynamics and the evolution of genetic drug resistance in our study.

To model partial drug resistance due to gene-expression noise, it was assumed that *k*_*N*_ *≤k*_*G*_, with genetic drug resistance mutations providing the greatest level of resistance. Genetic mutations were also assumed to be permanent in our simulations.

The switching rates between *S* and *N* are based on experimental estimates [3] and ranged between *r*_*S,N*_ = 0.0035 /hr and *r*_*N,S*_ = 0.0625 /hr. The mutation rate from *N* to *G* was based on a previous modeling-experimental study [2], which ranged from 10^− 6^ to 10^− 7^ per cell division, and was assigned a value of *r*_*G,N*_ = 0.3×10^− 6^ /hr in our simulations. The small magnitude of *r*_*G,N*_ relative to the growth and switching rate parameters was found to cause negligible differences in our simulation results.

The death rates *δ*_*S*_ and *δ*_*N*_ in cidal drug environments were scanned over similar order of magnitudes as the growth rates. To satisfy the assumption that the *N* subpopulation is more fit than the *S* subpopulation, we considered constant death rates given by:

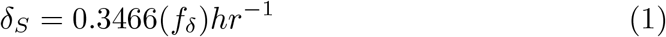

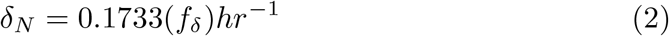

where *f*_*δ*_ is a scaling constant. To model low-strength, intermediate-strength, and high-strength cidal drug treatments we performed simulations for *f*_*δ*_ = 1, *f*_*δ*_ = 2, and *f*_*δ*_ = 4, respectively.

The deterministic and stochastic population models provide a means to simulate the effects of non-genetic resistance on the evolution of drug resistance. The effects of static drugs (which reduce cell growth but do not kill cells) were modelled by arresting the growth of *S* (*k*_*S*_ = 0) and reducing the growth rates of *N* and *G* (*k*_*N*_ and *k*_*G*_, respectively) and by setting the death rates to zero (*δ* _*S*_ = *δ* _*N*_ = *δ* _*G*_ = 0). Cidal drugs (which eventually kill all non-genetically resistant cells) were modelled by non-zero death rates for *S* and *N* (*δ* _*S*_ and *δ* _*N*_, respectively). To model the partial drug resistance of *N, δ* _*N*_ *< δ* _*S*_.

The parameter values given in Table S1 were also used for the stochastic simulations, as all the corresponding reactions in the stochastic population model were of zeroth-order or first-order [5]. Note that in the stochastic population model the values of *S*_*i*_, *N*_*i*_, and *G*_*i*_ are exact numbers of cells (*S*_*i*_ = 5.5×10^5^ cells and *N*_*i*_ = 5.5×10^4^ cells), which we used as the initial conditions to investigate the stochastic transition from non-genetic to genetic resistance as the number of cells in the population approached zero (extinction) due to cidal drug treatment. The *S* to *N* (and vice-versa) phenotype switching rates (*r*_*N,S*_ and *r*_*S,N*_, respectively), the *N* to *G* mutation rate (*r*_*G,N*_), and the growth and death rates (*k*_*i*_ and *δ* _*i*_, respectively, where *i ∈{S, N, G}*) are all given as probability per unit time in the stochastic population model.

### 2 Low-Strength Cidal Drug Condition

For the low-strength cidal drug environment, we considered a cidal death rate scaling factor of *f*_*δ*_ = 1.

Subpopulation trajectories, the fraction of genetically drug resistant cells in the population *G/T*, and the population rate of change *dT/dt* for the low-strength cidal drug scenario are shown in Figure S7.

A fixation time heat map for this case is shown in Figure S9A.

### 3 High-Strength Cidal Drug Condition

For the high-strength cidal drug environment, we considered a cidal death rate scaling factor of *f*_*δ*_ = 4.

Subpopulation trajectories, the fraction of genetically drug resistant cells in the population *G/T*, and the population rate of change *dT/dt* for the high-strength cidal drug scenario are shown in Figure S8.

A fixation time heat map for this case is shown in Figure S9B.

### 4 Survival and Extinction Scenarios

Representative survival and extinction trajectories from the stochastic simulations are shown in Figure S14. Figure S14A shows a case where the cell population survives cidal drug treatment due to *G* emerging before *S* and *N* go extinct. *G* subsequently takes over the population in this scenario. In contrast, Figure S14B shows a case where the cell population goes extinct due to *S* and *N* reaching zero cells before *G* appears.

### 5 Parameter Scans

To show the applicability of the mathematical model to other model organisms, simulation results for order-of-magnitude parameter scans of the switching rates *r*_*N,S*_ and *r*_*S,N*_, the mutation rate *r*_*G,N*_, and the growth rates *k*_*N*_ and *k*_*G*_ for the static (Figs. S2-S5) and cidal (*f*_*δ*_ = 2; Figs. S9-S13) drug scenario considered in the main text. These figures demonstrate that the conclusions made in the main text hold for a wide range of parameters.

### 6 Analytic Steady-State Solution

The equations in the mathematical model (Eqs. (1)-(3) in the main text) were set equal to zero and solved algebraically to yield the following steady-state solutions for the susceptible (*S*^*ss*^), non-genetically resistant (*N*^*ss*^), and genetically resistant (*G*^*ss*^) subpopulations:

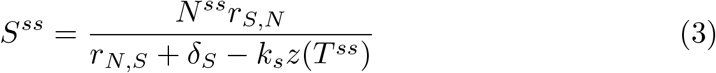

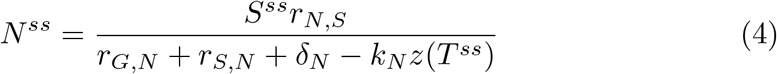

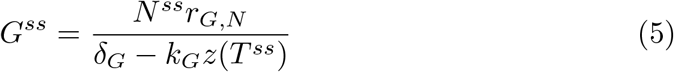

where *T* ^*ss*^ is the steady-state concentration of the entire population (i.e., *T* ^*ss*^ = *S*^*ss*^ + *N*^*ss*^ + *G*^*ss*^), *r*_*S,N*_ is the switching rate from *N* to *S, r*_*N,S*_ is the switching rate from *S* to *N, r*_*G,N*_ is the mutation rate from *N* to *G*, and *δ* _*S*_, and *δ* _*N*_, *δ* _*G*_ are the death rates of *S, N*, and *G*, respectively. Note that here 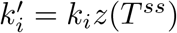, where z(*T* ^*ss*^) the competition function at steady-state and is otherwise similar to the competition function described by Eq. (6) in the main text. The following conditions must be satisfied for *N*^*ss*^ and *G*^*ss*^ to be non-negative and non-infinite:

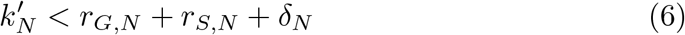

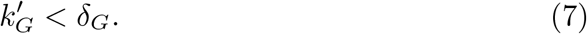

### 7 Partially Drug-Resistant Genetic Mutant

Figure S6 shows a case where *G* is assigned a non-zero death rate and hence is not fully resistant to the cidal drug. For this case, *f*_*δ*_ = 1 and *G* is assigned a death rate of *δ* _*G*_ = 0.2600(*f*_*δ*_) /hr. The time series show three different cases. The first case is where *N* and *G* have the same low-fitness growth rate, resulting in the total population moving towards extinction over time (solid blue lines in Fig. S6). The second case captures a scenario where *N* and *G* again have the same growth rate but now have intermediate fitness during drug treatment; in this case, all subpopulations reach a non-zero steady-state with *S*^*SS*^ *>> G*^*SS*^ and *N*^*SS*^ *>> G*^*SS*^ (dash-dotted red lines in Fig. S6). The last case shows a situation where *N* has a growth rate that is greater than the growth rate of *G*, which also results in non-zero subpopulation steady states (dashed yellow lines in Fig. S6). Other cases were investigated and were found to show similar trends of either full population extinction or non-zero subpopulation steady states, depending on the relative fitness and the susceptibility to drug treatment (data not shown).

## Supporting Table

**Table 1:**
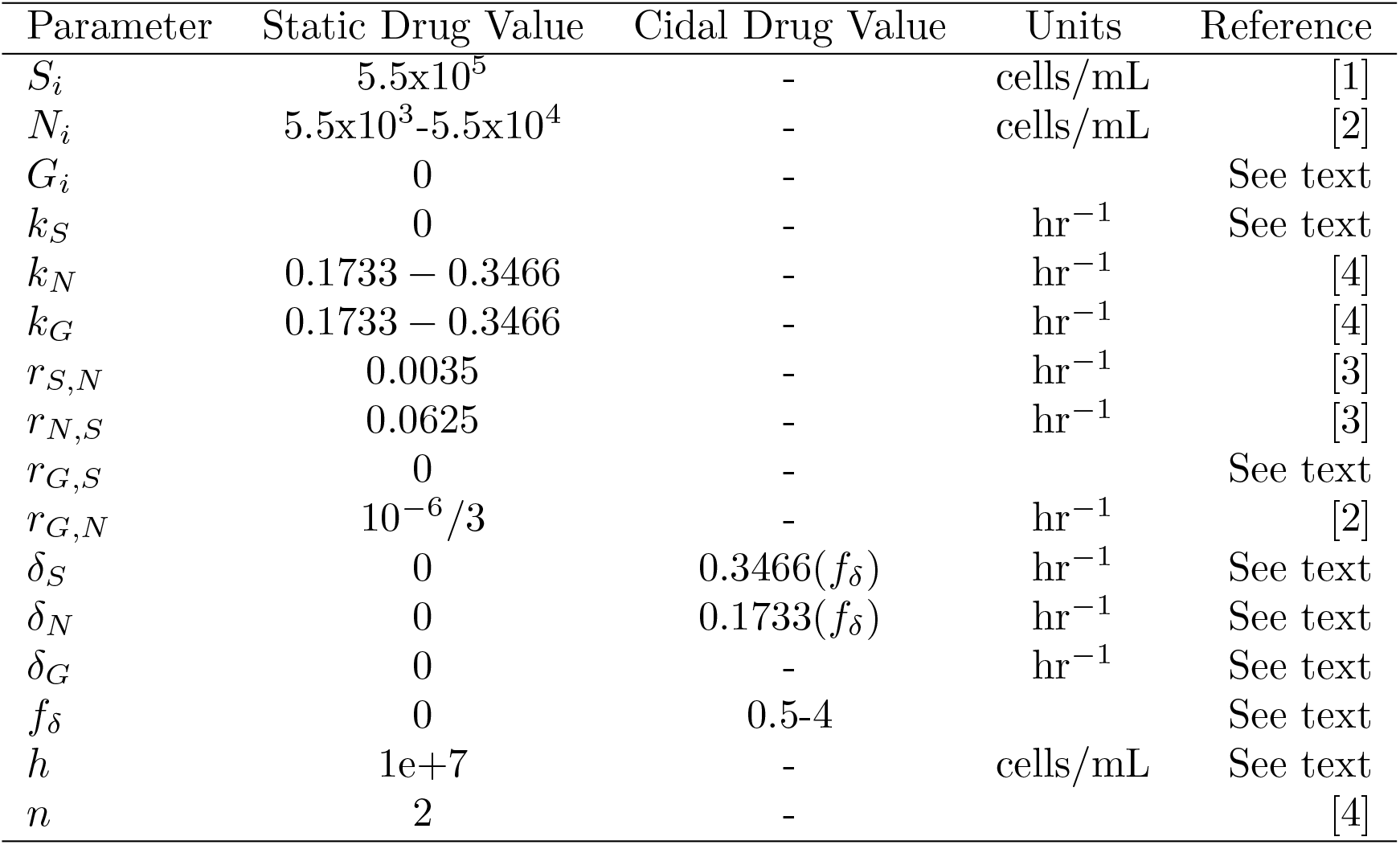
Parameters used for numerically simulating the deterministic population dynamics model in static and cidal drug conditions. A horizontal line in the ‘Cidal Drug Value’ column indicates that the value is the same as the corresponding value in the ‘Static Drug Value’ column. A blank entry in the ‘Units’ column indicates no units and ‘See text’ in the ‘Reference’ indicates that the justification for the parameter value is giving within the text of the main text or of the Supporting Material.

## Supporting Figures

**Figure 1:**
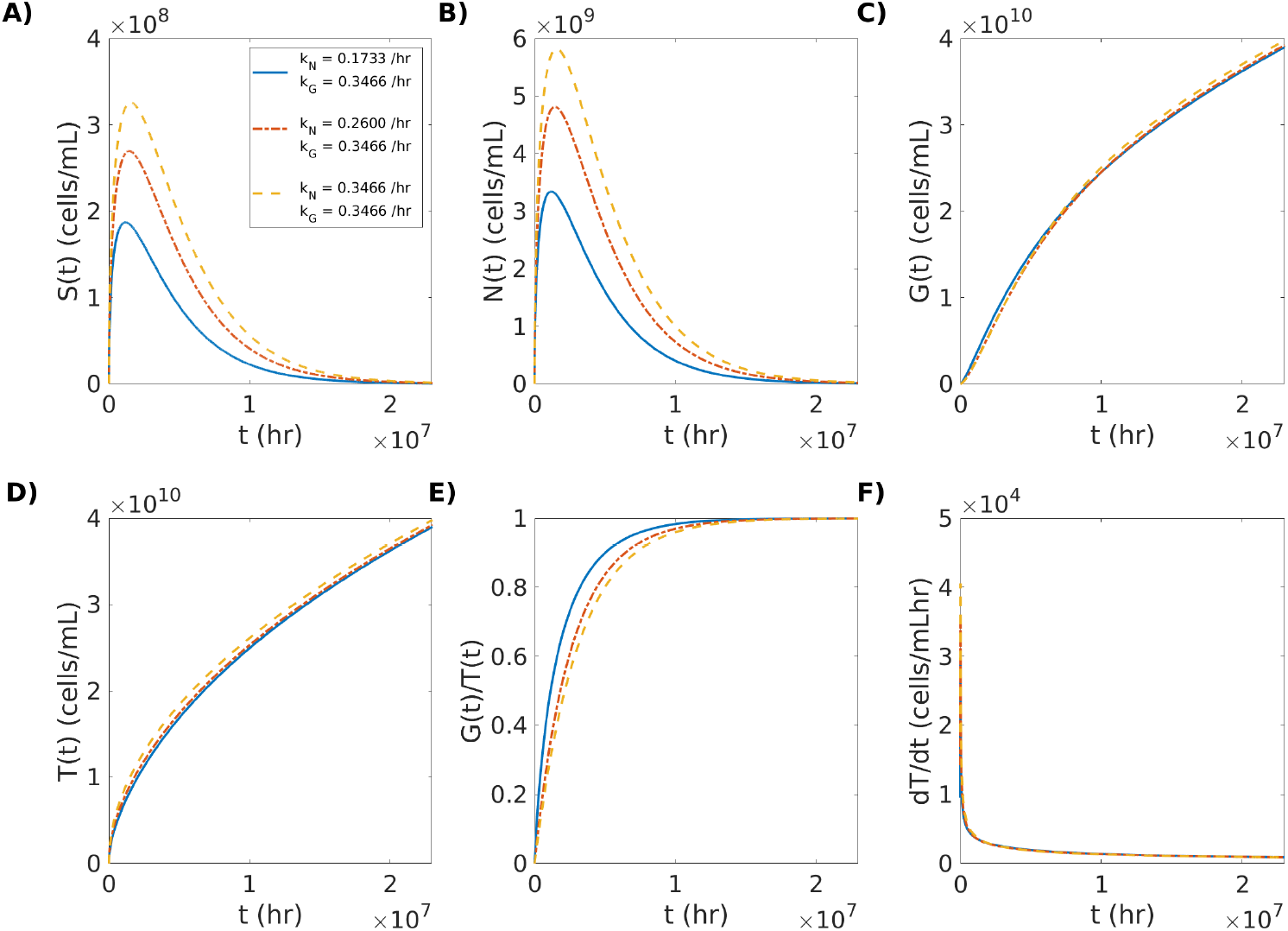
Resource competition drives drug-resistant subpopulations extinct over long time scales during static drug treatment. (A) The growth curve of the drug susceptible (*S*) subpopulation. (B) The growth curve of the non-genetically drug resistant (*N*) subpopulation. (C) The growth curve of the genetically drug resistant (*G*) subpopulation. (D) The growth curve of the total population (*T*). (E) The fraction of *G*(*t*) in *T*. (F) The rate of change in the size of *T* (*dT/dt*) as a function of time. Each colored line represents a different numerical simulation using the growth values shown in the legend in (A), with the solid blue line representing the lowest level of *N* fitness (0.1733 /hr), the red dash-dotted line an intermediate level of *N* fitness (0.2600 /hr), and the yellow dashed line the highest level *N* fitness (0.3466 /hr), relative to the fitness of *G* (0.3466 /hr).

**Figure 2:**
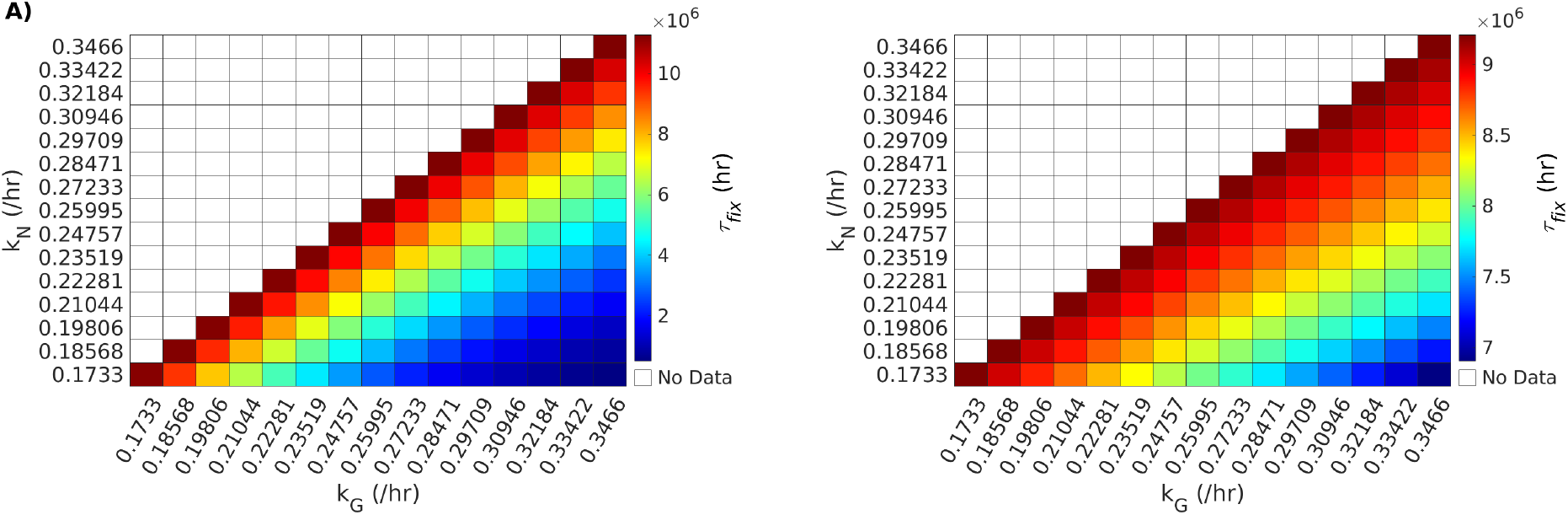
Genetic fixation results in static drug environment for *r*_*N,S*_ = 0:001 /hr (A) and *r*_*N,S*_ = 0:1 /hr (B).

**Figure 3:**
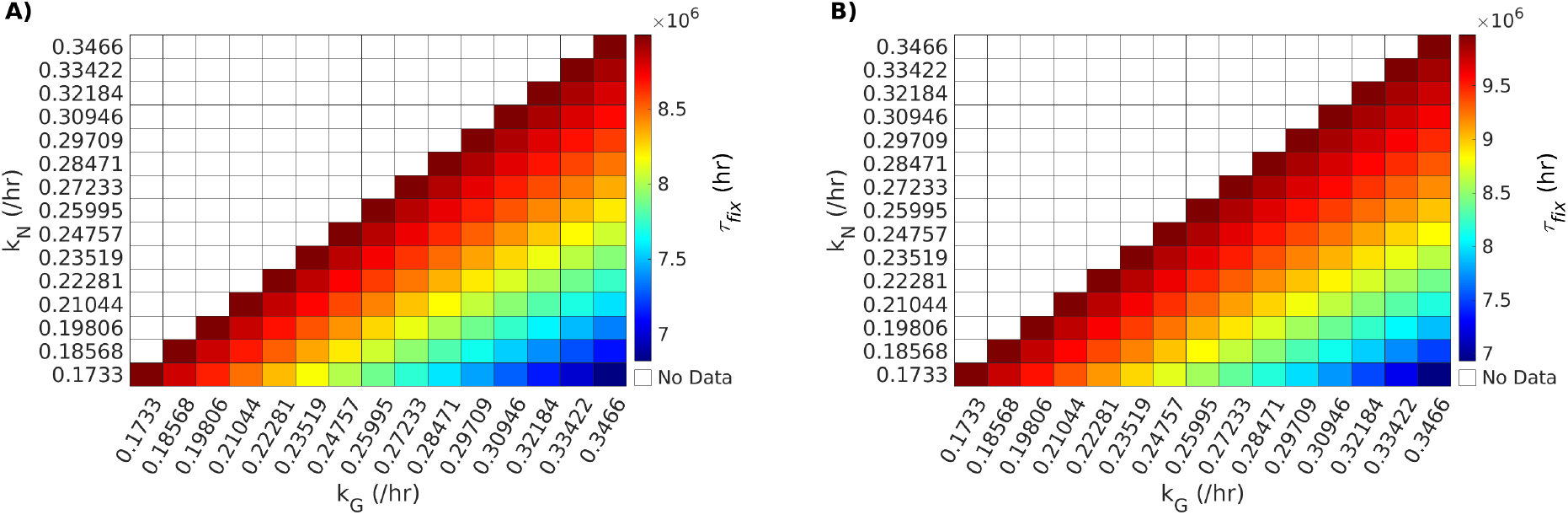
Genetic fixation results in static drug environment for *r*_*N,S*_ = 0:001 /hr (A) and *r*_*N,S*_ = 0:1 /hr (B).

**Figure 4:**
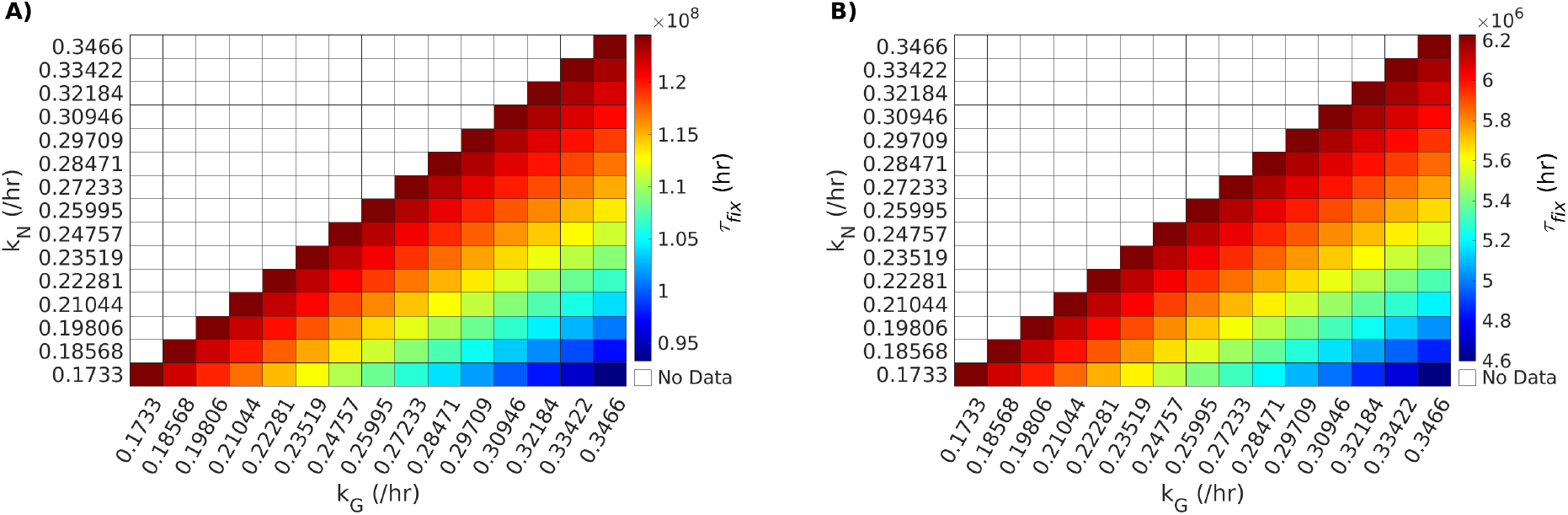
Genetic fixation results in static drug environment for *r*_*G,N*_ = 10^− 7^ /4hr (A) and *r*_*G,N*_ = 10^− 6^ /2hr (B).

**Figure 5:**
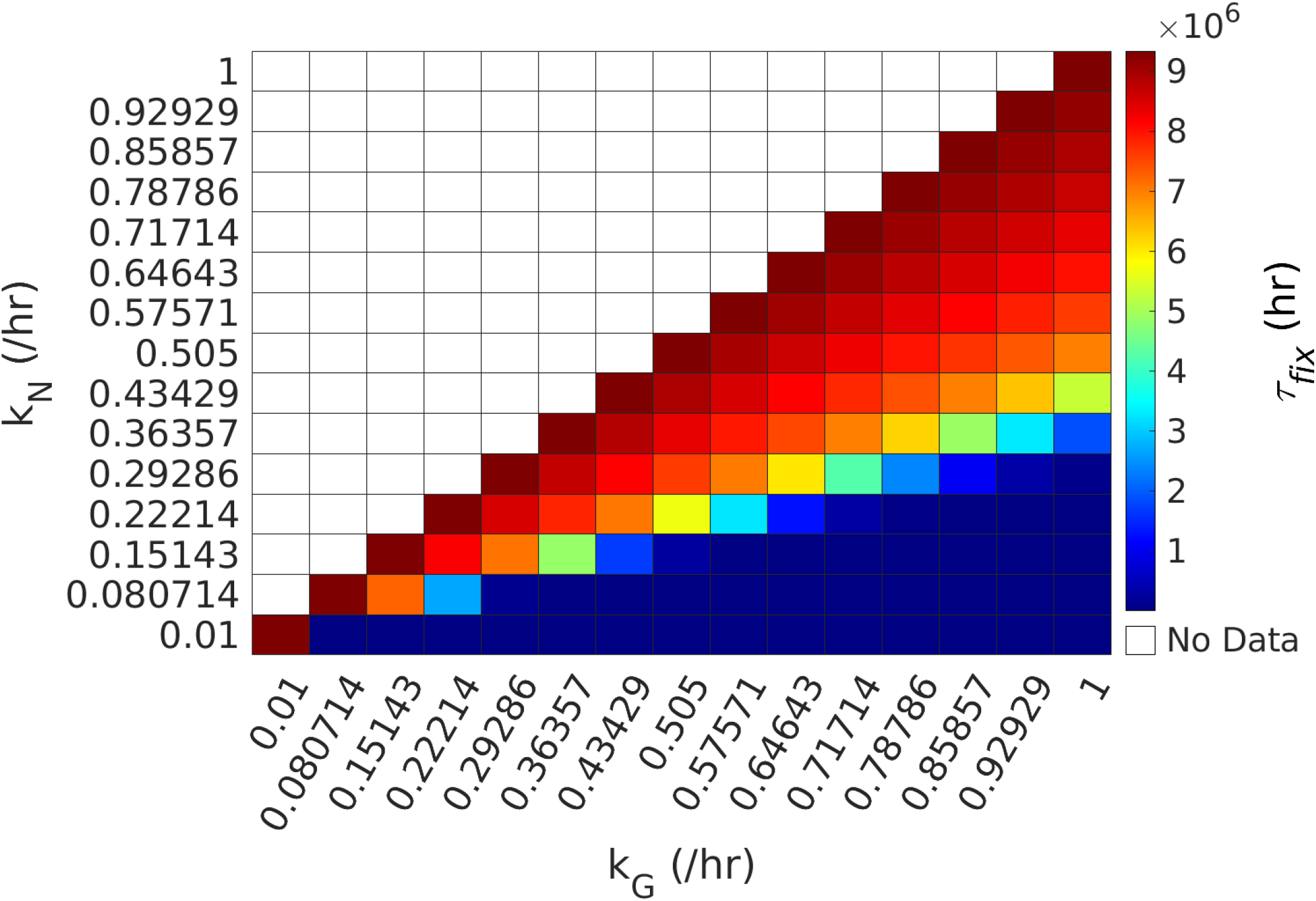
Genetic fixation results in static drug environment for growth rates *k*_*N*_ and *k*_*G*_ scanned from 0.01 /hr to 1 /hr. Note that the large range of values (ranging from 10^2^ hr to 10^6^ hr for high *k*_*G*_ and low *k*_*N*_) resulted in the same color bins in the MATLAB heatmap, though at a finer resolution (not shown) the trends were qualitatively the same as those presented in the main text.

**Figure 6:**
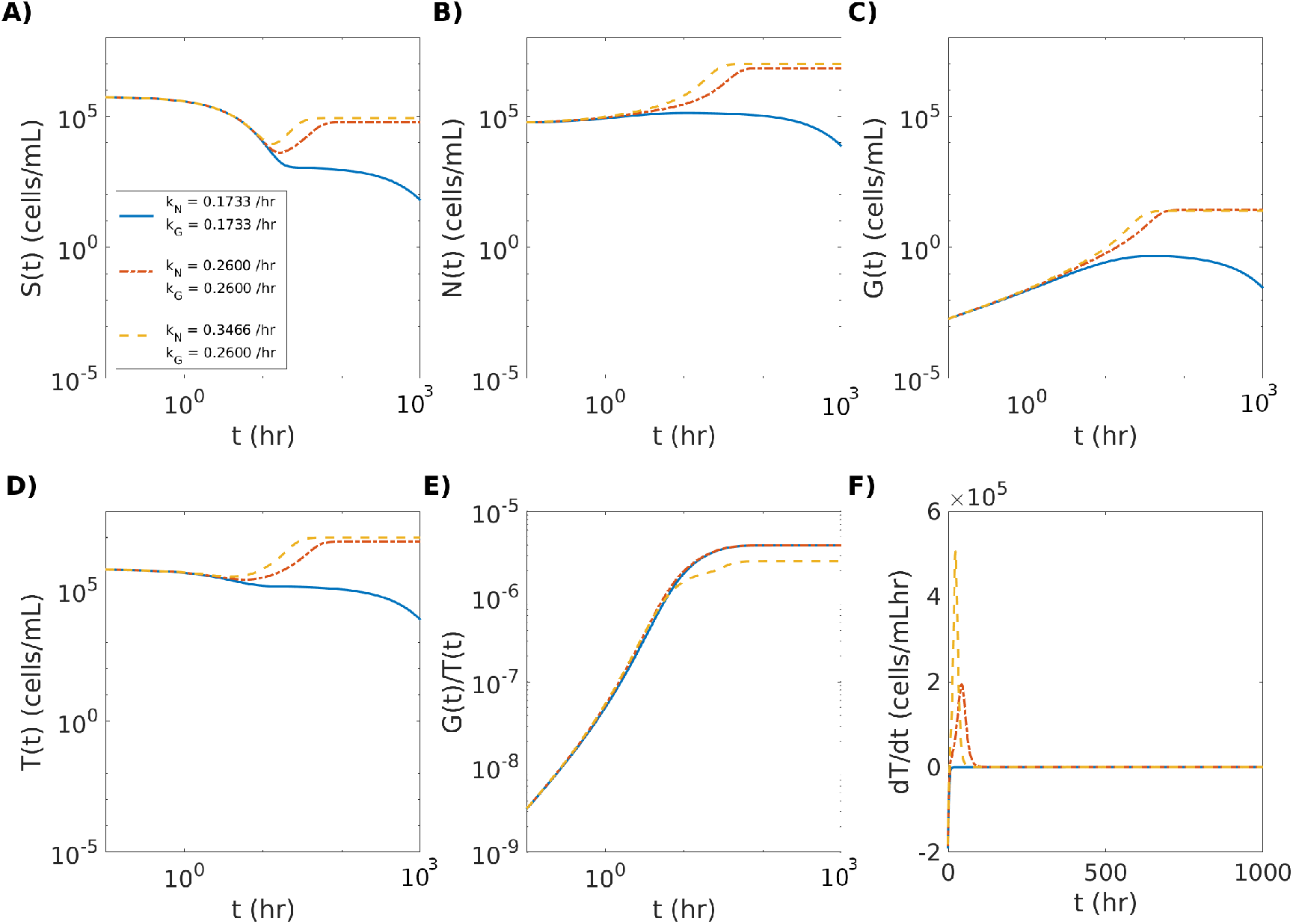
Partially resistant genetic mutation results in either total population extinction or non-zero steady-state population size during cidal drug treatment (*f*_*δ*_ = 1). (A) The growth curve of the fully drug susceptible (*S*) subpopulation. (B) The growth curve of the non-genetically drug resistant (*N*) subpopulation. (C) The growth curve of the genetically partially drug resistant (*G*) subpopulation. (D) The growth curve of the total population (*T*). (E) The fraction of *G*(*t*) in *T*. (F) The rate of change in the size of *T* (*dT/dt*) as a function of time. Each colored line represents a different numerical simulation using the growth values shown in the legend in (A), with the solid blue line representing the lowest level of *N* and *G* fitness (0.1733 /hr), the red dash-dotted line an intermediate level of *N* and *G* fitness (0.2600 /hr), and the yellow dashed line the highest level *N* fitness (0.3466 /hr), relative to the intermediate fitness level of *G* (0.2600 /hr).

**Figure 7:**
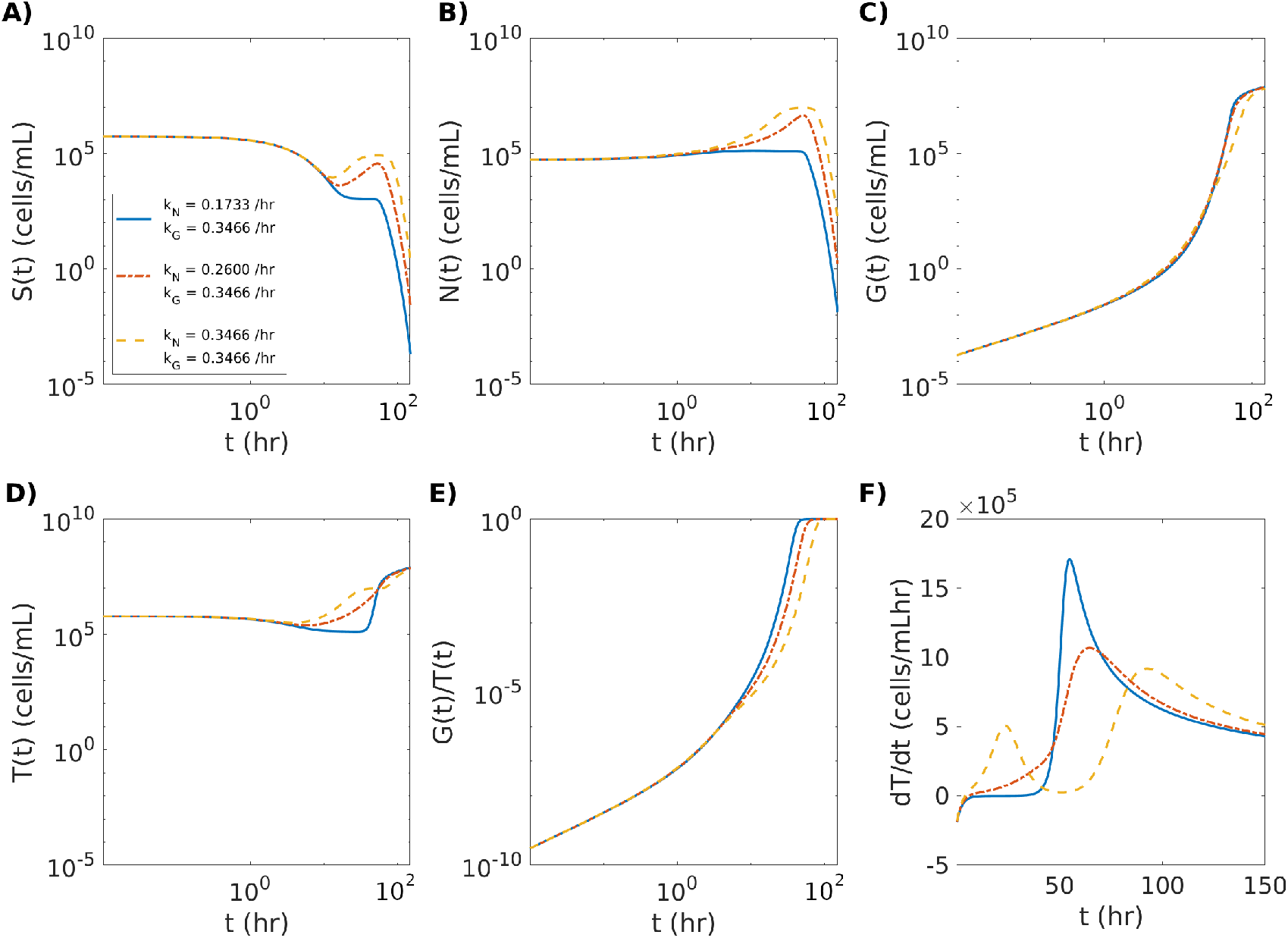
Growth of the genetically resistant subpopulation is hindered by the growth of the non-genetically resistant subpopulation in a low-strength cidal drug environment. (A) The growth curve of the drug susceptible (*S*) subpopulation. (B) The growth curve of the non-genetically drug resistant (*N*) subpopulation. (C) The growth curve of the genetically drug resistant (*G*) subpopulation. (D) The growth curve of the total population (*T*). (E) The fraction of *G*(*t*) in the total population *T*. (F) The rate of change in the size of *T* (*dT/dt*) as a function of time. Each colored line represents a different numerical simulation using the growth values shown in the legend in (A), with the solid blue line representing the lowest level of *N* fitness (0.1733 /hr), the red dash-dotted line an intermediate level of *N* fitness (0.2600/hr), and the yellow dashed line the highest level *N* fitness (0.3466 /hr), relative to the fitness of *G* (0.3466 /hr) under low-strength cidal drug exposure. The cidal death rate scaling factor was set to *f*_*δ*_ = 1.

**Figure 8:**
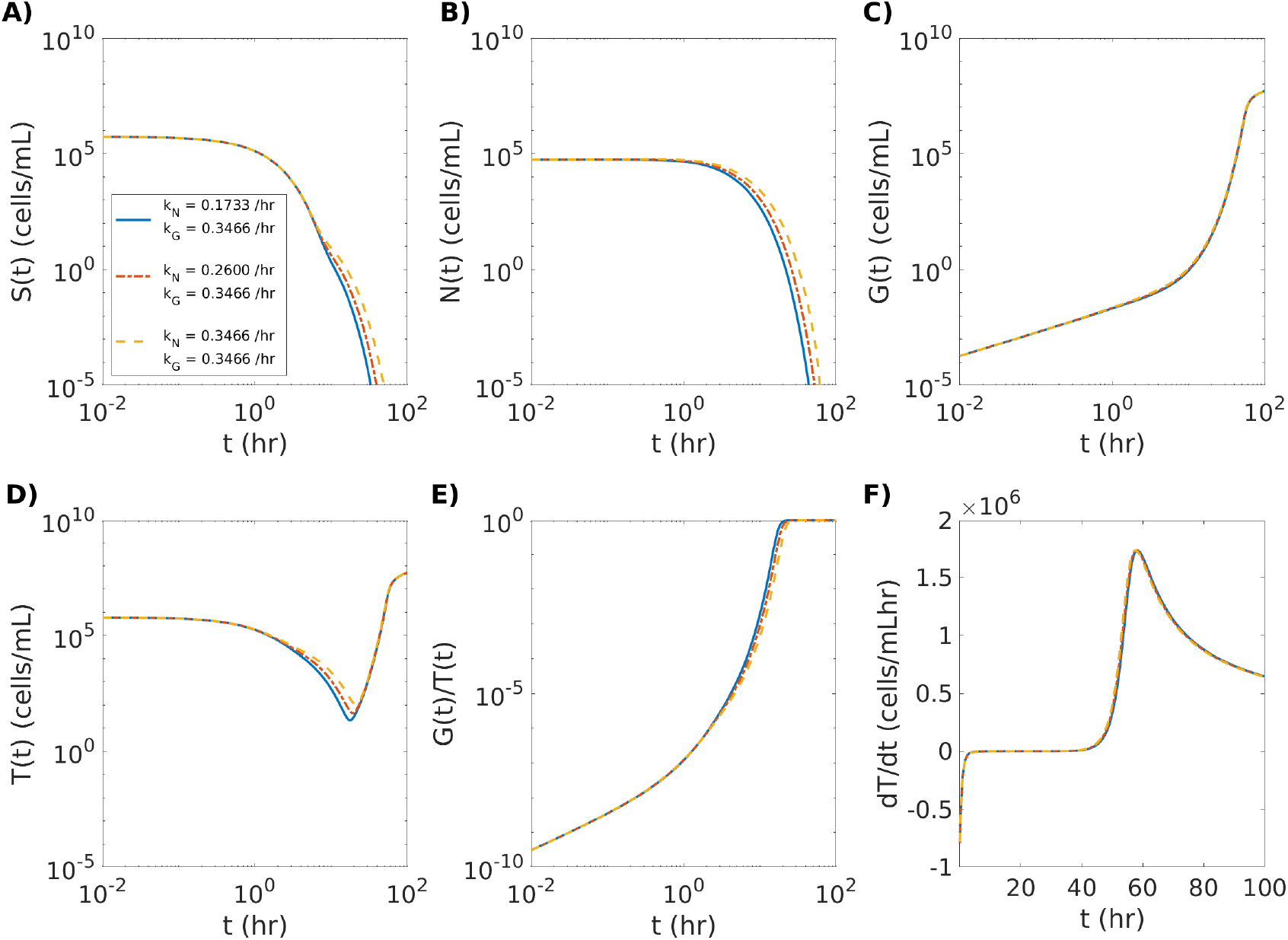
Growth of the genetically resistant subpopulation is hindered by the growth of the non-genetically resistant subpopulation in a high-strength cidal drug environment. (A) The growth curve of the drug susceptible (*S*) subpopulation. (B) The growth curve of the non-genetically drug resistant (*N*) subpopulation. (C) The growth curve of the genetically drug resistant (*G*) subpopulation. (D) The growth curve of the total population (*T*). (E) The fraction of *G*(*t*) in the total population *T*. (F) The rate of change in the size of *T* (*dT/dt*) as a function of time. Each colored line represents a different numerical simulation using the growth values shown in the legend in (A), with the solid blue line representing the lowest level of *N* fitness (0.1733 /hr), the red dash-dotted line an intermediate level of *N* fitness (0.2600 /hr), and the yellow dashed line the highest level *N* fitness (0.3466 /hr), relative to the fitness of *G* (0.3466 /hr) under high-strength cidal drug exposure. The cidal death rate scaling factor was set to *f*_*δ*_ = 4.

**Figure 9:**
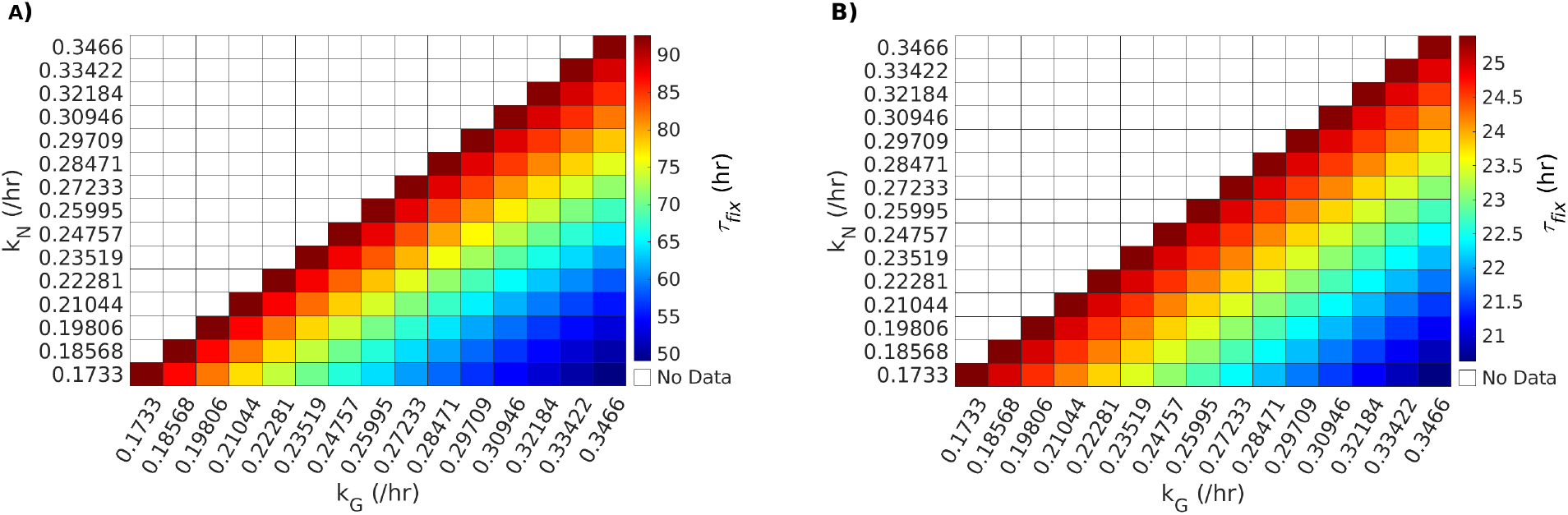
Drug resistance of the non-genetic subpopulation slows the development of the genetically drug resistant subpopulation in a cidal drug environment. (A) Heat map shows the effect of the growth rates of the non-genetically resistant (*k*_*N*_) and genetically resistant (*k*_*G*_) subpopulations on the fixation time (*τ*_*fix*_) of *G* under low-strength cidal drug treatment. The cidal death rate scaling factor was set to *f*_*δ*_ = 1 for low-strength cidal drug scenario. (B) Heat map shows the effect of *k*_*N*_ and *k*_*G*_ on the fixation time (*τ*_*fix*_) of *G* under high-strength cidal drug treatment. The cidal death rate scaling factor was set to *f*_*δ*_ = 4 for the high-strength cidal drug scenario. In the heat maps shown in (A) and (B), each bin corresponds to a simulation for a combination of *k*_*N*_ and *k*_*G*_ parameter values. The color maps show the fixation time in hours. As *k*_*N*_ is less than or equal to *k*_*G*_ in our model, numerical simulation data does not appear in the upper diagonal of the heat maps.

**Figure 10:**
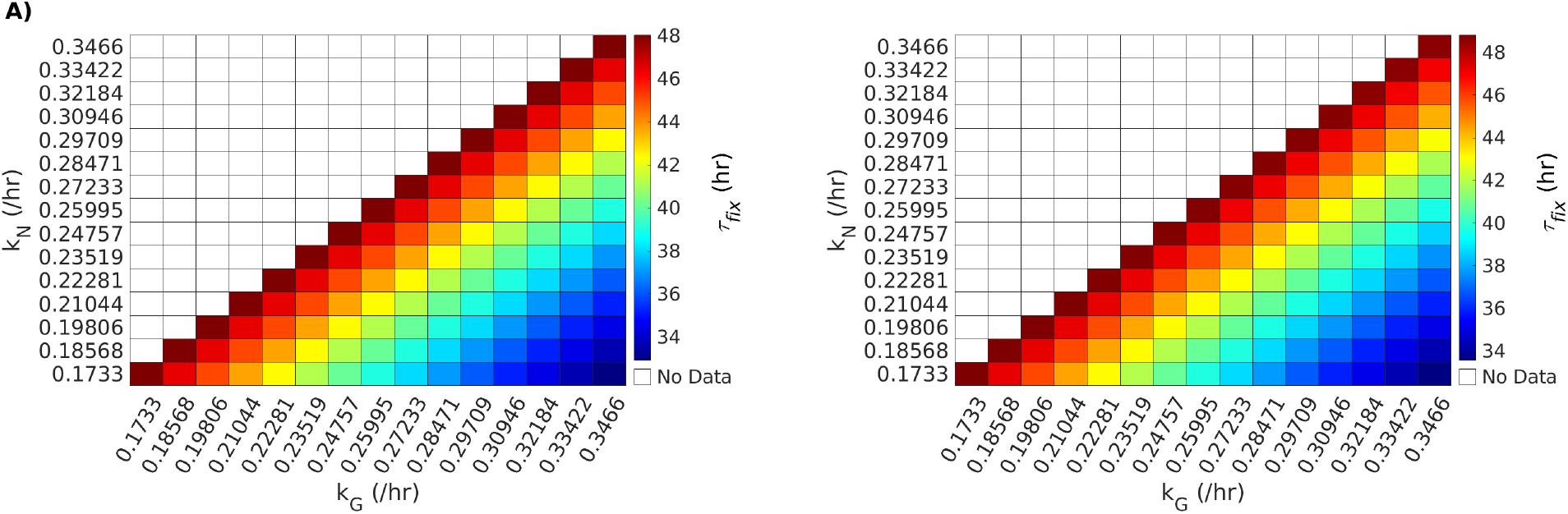
Genetic fixation results in cidal drug environment (*δ* _*f*_ = 2) for *r*_*N,S*_ = 0.001 /hr (A) and *r*_*N,S*_ = 0.1 /hr (B).

**Figure 11:**
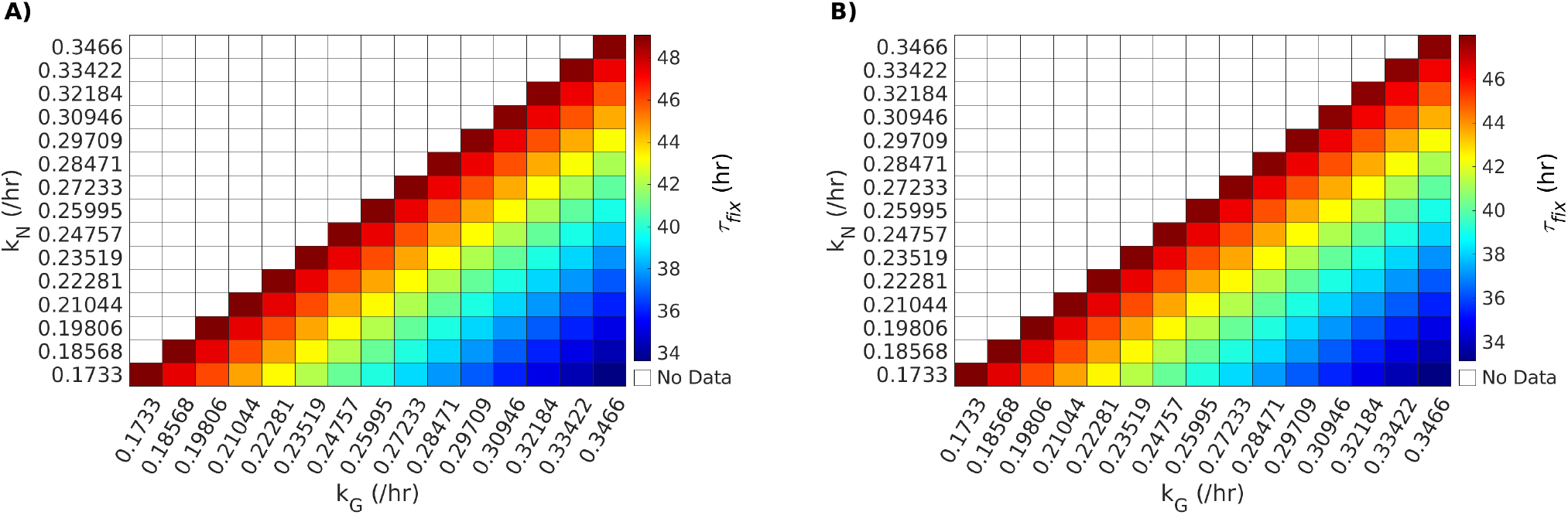
Genetic fixation results in cidal drug environment (*δ* _*f*_ = 2) for *r*_*S,N*_ = 0.0001 /hr (A) and *r*_*S,N*_ = 0.01 /hr (B).

**Figure 12:**
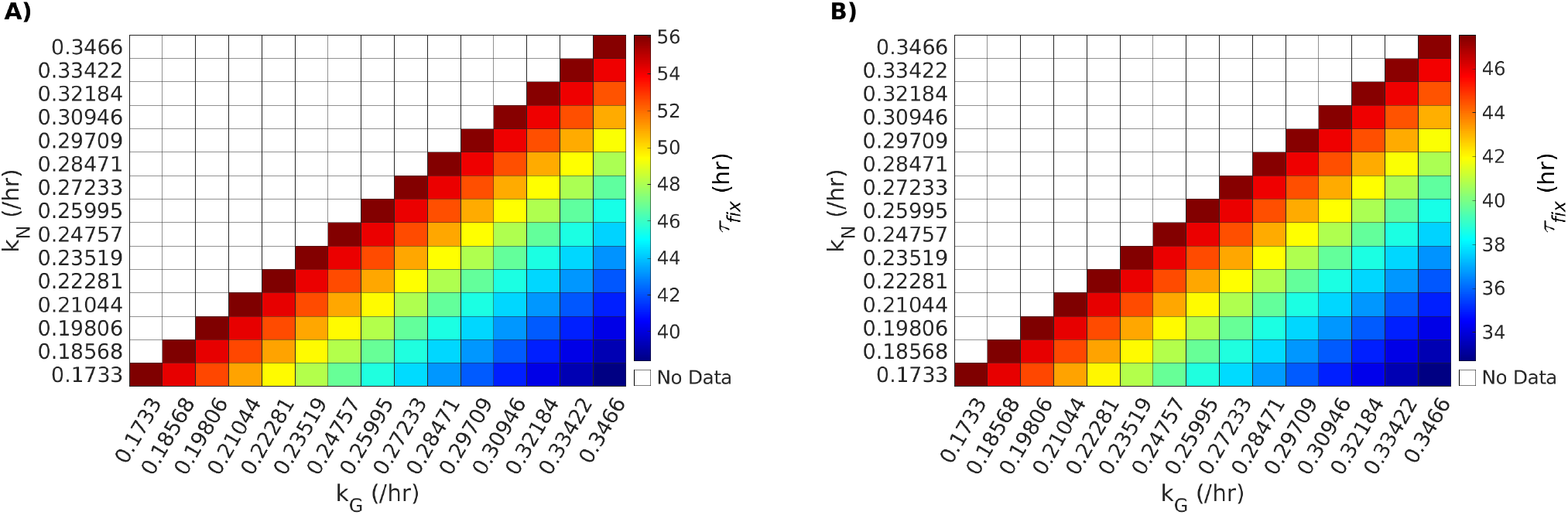
Genetic fixation results in cidal drug environment (*δ* _*f*_ = 2) for *r*_*G,N*_ = 10^− 7^ /4hr (A) and *r*_*G,N*_ = 10^− 6^ /2hr (B).

**Figure 13:**
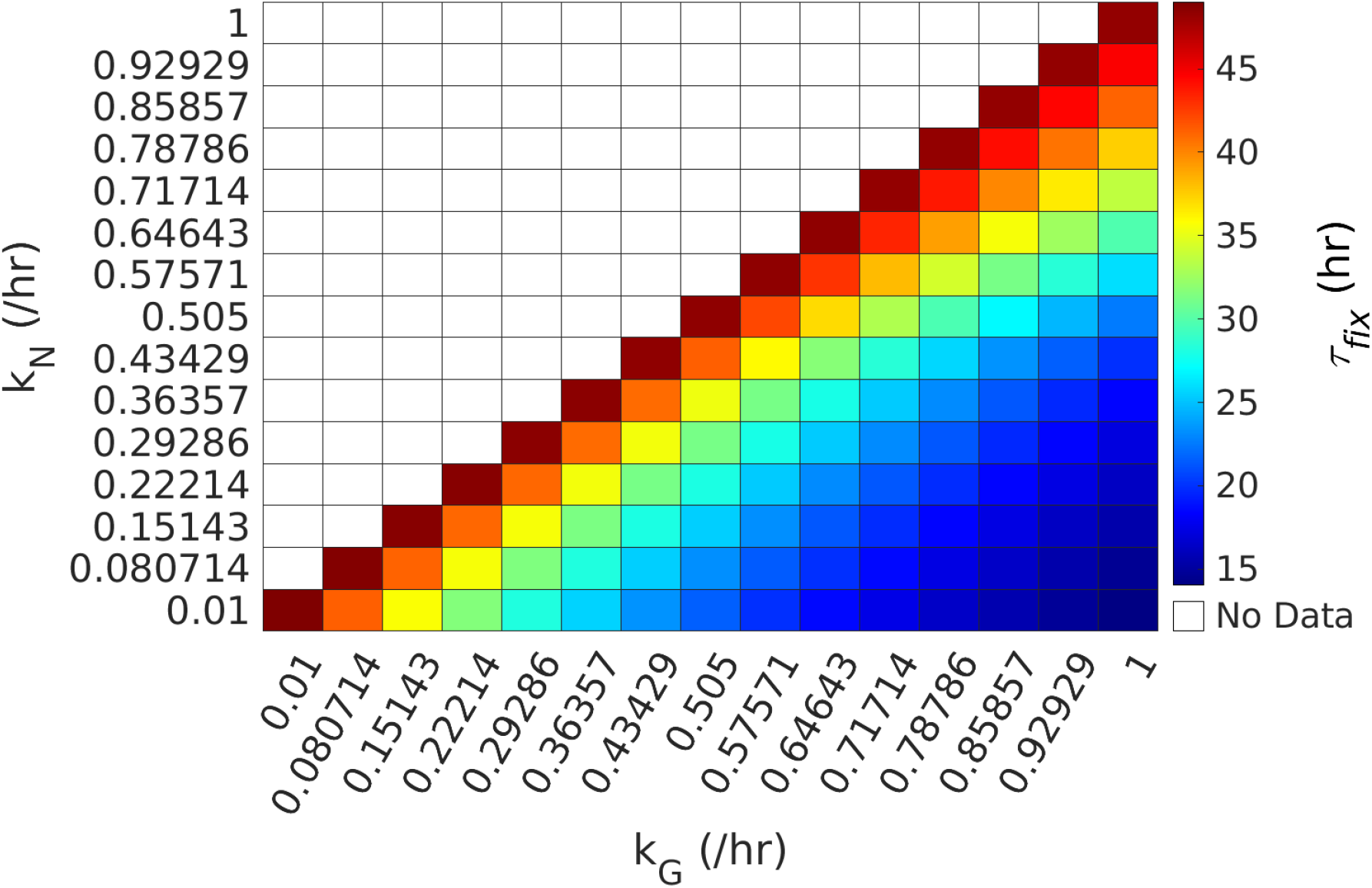
Genetic fixation results in cidal drug environment (*δ* _*f*_ = 2) for growth rates *k*_*N*_ and *k*_*G*_ scanned from 0.01 /hr to 1 /hr.

**Figure 14:**
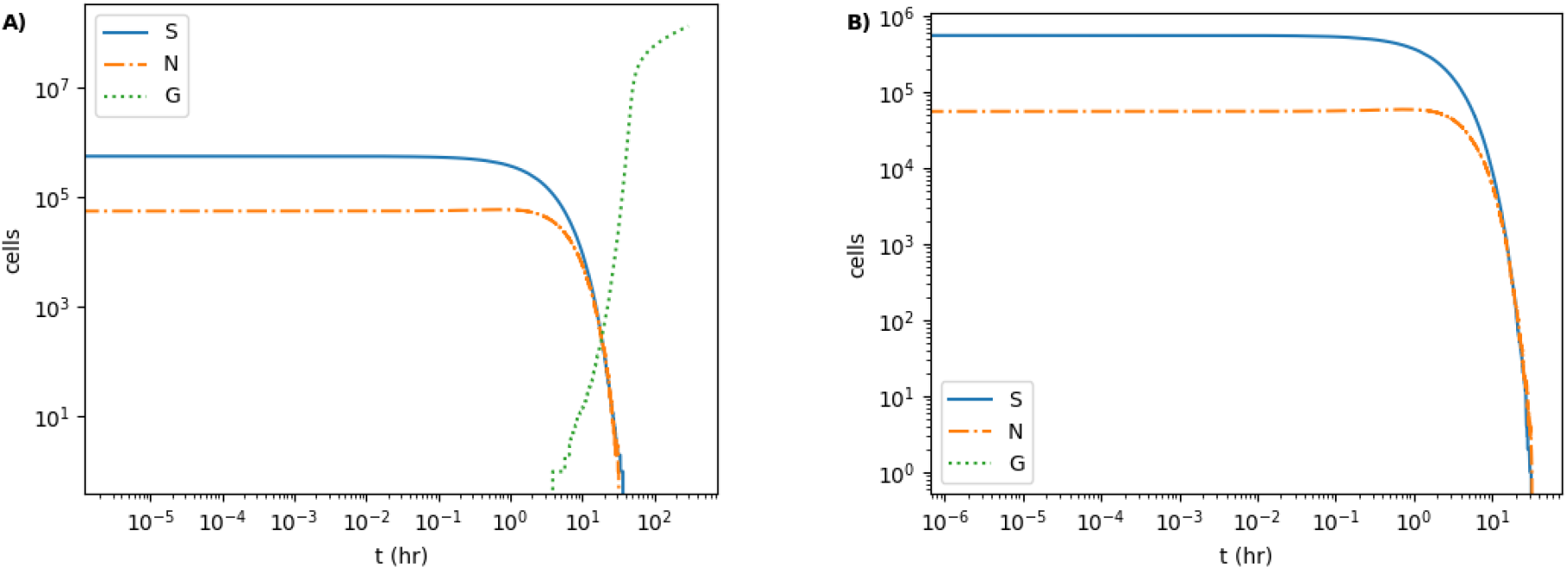
Cell population survival and extinction during cidal drug treatment. (A) A representative population non-extinction case where *G* appears before *S* and *N* go extinct. (B) A representative population extinction case where *S* and *N* go extinct before *G* appears in the population. For (A) and (B), the parameter values *k*_*N*_ = 0.1733 /hr and *f*_*δ*_ = 2 were used in the stochastic simulations.

## Notes

### Competing Interest Statement

The authors have declared no competing interest.

